# Reassessing the helix bundle crossing model for gating in a non-selective ion channel

**DOI:** 10.1101/2022.04.18.488652

**Authors:** Vilius Kurauskas, Marco Tonelli, Katherine Henzler-Wildman

## Abstract

A critical part of ion channel function is the ability to open and close in response to stimuli, and thus conduct ions in a regulated fashion. While X-ray diffraction studies of ion channels suggested a general steric gating mechanism located at the helix bundle crossing (HBC), recent functional studies on several channels indicate that the helix bundle crossing is open even in closed, non-conductive channels. Two NaK channel variants were crystallized in very different, open and closed conformations and served as an important model of the HBC gating hypothesis. However, neither of these NaK variants are conductive in liposomes unless phenylalanine 92 is mutated to alanine (F92A). Here we use NMR to probe distances at near-atomic resolution of the two NaK variants in lipid bicelles. We demonstrate that in contrast to the crystal structures, both NaK variants are in a fully open conformation, akin to the well known MthK channel structure were the HBC is widely open. Further inquiry into the gating mechanism suggests that the selectivity filter and pore helix are coupled to the M_2_ helix below and undergo changes in structure when F92 is mutated. Overall, our data shows that NaK exhibits coupling between the selectivity filter and HBC similar to K^+^ channels and has a more complex gating mechanism than previously thought.

## INTRODUCTION

Ion channel function is defined by the selectivity of the channel and the gating mechanism. Central to understanding the function is the knowledge of ion channel structure. The first X-ray crystal structures of potassium channels were pivotal in cementing the image of widening of the helix bundle crossing (HBC) from a sterically occluded state to a more open state as the principal determinant that initiates ionic current. For example, the initial structure of KcsA has a bundle crossing too narrow for K^+^ ions to pass ^1^ which was suggested to be a deactivated nonconductive conformation, while the subsequent structure of MthK has a widely open bundle ^2^, that was interpreted as open, conductive channel. Over the years X-ray and EM structures of dozens of different channels have appeared bound to various activators and inhibitors, but almost none of the channels clearly displayed both of the HBC conformations expected for open and closed states ^3^. Out of the small handful of examples where mutagenesis was used to obtain atomic resolution structures of both conformations ^4–6^, the structures of NaK channel from *B. cereus* appears to be the clearest example of gating at the HBC. The full-length NaK X-ray structure displays a narrowly constricted bundle-crossing ^7^, while the removal of an amphiphatic helix (M_0_ removed, NaKΔ19) renders this channel widely open ^8^ – nicely fitting into the narrative of steric gating at the HBC.

Such a model of gating at the HBC, as derived from observations in NaK and other model channels, has strongly affected how structure-function experiments of ion channels were designed and interpreted. Putative elusive conformations of KcsA were sought out by cross-linking M_2_ helices with disulfide bonds ^9^ or mutating residues to stabilize the closed HBC conformation ^6^. Meanwhile, solid and solution-state NMR studies on KcsA and KirBac1.1 have interpreted chemical shift perturbation data as supporting the two state HBC gating model ^10–15^. However, in the case of KirBac3.1 it was later demonstrated that inner helix bundle stabilized with disulfide bonds in a closed conformation still allows K^+^ ion conduction ^16^ and NMR chemical shift guided structural investigation of the conductive conformation of KcsA suggested a much narrower HBC opening than initially believed from the X-ray structures ^17^. The struggle of assigning structures with different HBC conformations to functional states continues with more recent EM structures, as exemplified by the TASK channel ^18,19^, where structural interpretations do not necessarily agree with the functional data and a residue far away from HBC appears to be critical for gating ^20^.

Even in the case of NaK, certain discrepancies remain regarding the connection between the putative open and closed X-ray structures of NaK and channel function. For example, Rb^+^ flux assays demonstrate only a marginal increase in flux for an ‘open’ NaKΔ19 ^21^ construct over the ‘closed’ fulllength channel, and an additional mutation, F92A is necessary ^22^ to obtain fluxes observable by single channel electrophysiology measurements. This observation made the authors themselves question whether the ‘open’ conformation of NaK is an artifact of crystal packing ^21^. In the case of KcsA the HBC is still fully open when the channel becomes non-conductive ^23^ and this inactivation is now believed to occur at the selectivity filter. As the degree of HBC opening correlates with this C-type inactivation, reconciling such confounding observations is necessary to understand ion channel function in general.

There are now examples in nearly all K^+^ channel families that are suggested to gate somewhere other than the classical HBC. Among them are MthK ^24–26^ and related eukaryotic BK channels ^27,28^, K_2_P channels, such as TREK ^20,29^, CNG channels MloK1 ^30^ and TAX-4 ^31^, and Kir channel KirBac3.1 ^16^. Even the seemingly strongly established model of KcsA opening at the HBC has been questioned ^3,32,33^. That many channels might not be principally gated at the HBC is further strengthened by the fact that a common class of activators can enter “closed” channels from the cytoplasmic side, including the Kv channel HERG, and activate them ^34^. In all of these cases, the main activation gate was instead proposed to be in the selectivity filter (SF), while the HBC was reduced to a secondary role in gating where change in the opening angle of the HBC helices transmits an allosteric activation signal ^35,36^ to the SF.

Investigation of how KcsA channels are gated at the selectivity filter illuminated the critical importance on an aromatic residue located in the middle of the pore lining M_2_ helix near the base of the SF, F103. It has been established that F103A attenuates allosteric coupling between the SF and C-terminal glutamates ^37^ in KcsA, and removal of this residue suppresses C-type inactivation ^23^. Interestingly, many K^+^ channels appear to be affected by substitutions in this region. Mutations in analogous position (TM2.6) to polar residues greatly increases open probability in all vertebrate K2P channels ^20^. Human Slo1 becomes active in the absence of Ca^2+^ after F380Y substitution ^38^ and A388C-MTSEA modification renders bovine CNGA1 conductive in the absence of ligand ^39^. Moreover, a substitution one residue downstream has been reported to alter K^+^ channel gating as well. Kv channel HERG harboring A653 substitutions no longer closes at physiologically relevant potentials and display changes in the rate of C-type inactivation ^40^ and polar mutations at this site makes MthK ^41^ consitutively active. In other cases, such as KCNQ3 (Kv7.3), this substitution (F344A/C/W) results in reduced flux ^42^. Therefore, residues just below the selectivity filter, especially residues corresponding to F103 and A104 in KcsA appear to be important in gating of a broad range of K^+^ channels.

To address the discrepancy between the NaK X-ray structures and its functional behavior, we investigated whether NaK is primarily gated at the HBC using solution NMR spectroscopy and solid-supported membrane electrophysiology (SSME) measurements. Unlike solid-state NMR ^43^ spectra of NaK in liposomes, solution state NMR spectra of NaK samples in lipid bicelles display clear resonances for residues throughout the M_2_ helix, including those near the HBC ^44^. Additionally, while substantial attention has been paid to investigating KcsA by both solution and solid-state NMR, these studies largely excluded detailed investigation of the HBC, only indirectly correlating chemical shift perturbation data with hypothesized changes in the structure ^12,14,17^: in part due to the ambiguity of the detailed atomic structures of different states ^17^. While a powerful technique for unraveling structure and dynamics in smaller proteins, for large molecular systems such as ion channels, obtaining the high quality NMR data necessary for meaningful interpretation can be a challenge. Here we use a combination of backbone and methyl resonances and the resolution provided by solution NMR to measure both chemical shift perturbation and site-specific distance restraints for NaK constructs in bicelles. As NaK presents two putative open and closed structures at atomic resolution, which can be easily obtained by truncation of M_0_ helix, this represents an ideal system for testing the competing hypotheses for the location of the ion channel gate by NMR.

## RESULTS

### Comparison of NaKΔ18 and NaK FL NMR “fingerprints”

In this work we use NaKΔ18, which is similar to the NaKΔ19 construct ^21^ used to characterize the structural and functional impact of removing the M_0_ helix, but leaves in place residue W18 (Figure 1a). This aromatic residue may help stabilize the channel in the membrane and improves the stability of the construct during purification (Fig. S1). We began by comparing the backbone amide chemical shifts of the putative “open” and “closed” channel constructs, NaKΔ18 ^44^ and full-length NaK (NaK FL), in lipid bicelles. Although we did not assign the backbone resonances in NaK FL spectra, assignments of more dispersed peaks can be easily transferred between the ^1^H^15^N-TROSY-HSQC spectra of the two variants (Fig SI2). While the overall pattern remains very similar, we observe a chemical shift perturbation (CSP) of 0.25ppm for the W18 indole nitrogen, which is of a similar magnitude as reported for W26 in KcsA ^11^ and 0.05-0.1ppm CSP for the residues on the M_2_ helix near the HBC. In general, the pattern of CSPs appears to be qualitatively consistent with the hypothesis of conformational change in the hinge region correlated to gating at the inner bundle crossing, although the magnitude of CSPs appears small for such a large conformational change.

**Figure 1.**
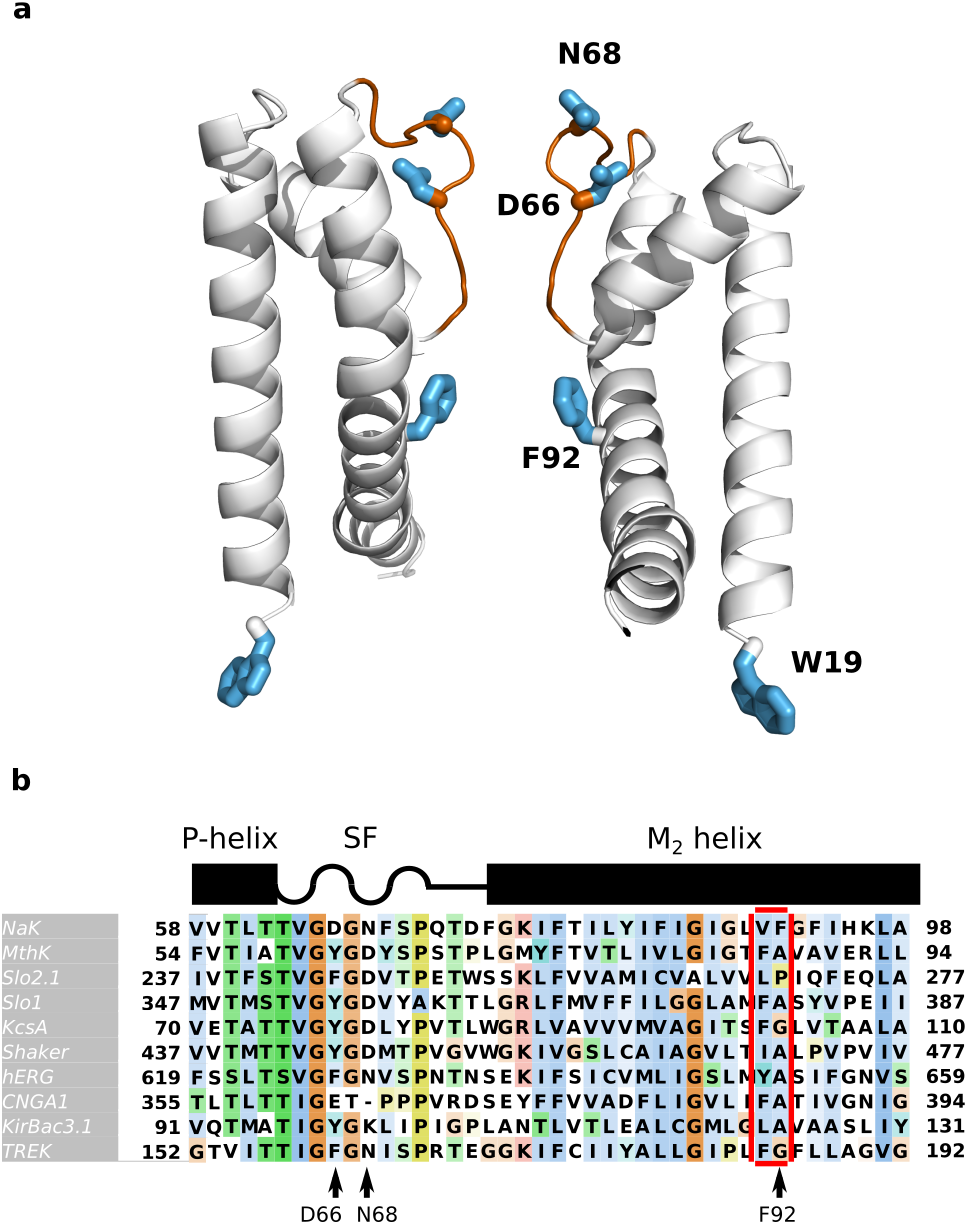
NaK structure and sequence alignment with related channels. a) Crystal structure of NaKΔ18 (3E8H) with only two opposing monomers displayed for clarity. The selectivity filter is colored in orange. Residues W19, F92, D66, and N68 are highlighted light blue sticks. b) Sequence alignment of ion channels with similar pore structure to KcsA. Black cartoon at top indicates the secondary structure elements. Residues V91-F92 are highlighted in red.

U-[^15^N^12^C^2^H] labeled NaKΔ18 in 100mM K^+^, displays good quality ^15^N-^1^H spectra at 40° C ^44^, but the correlation time in bicelle is ~45 ns for NaKΔ18 and even longer for NaK FL, which hampers a detailed structural comparison. We therefore used methyl groups as NMR observables, which have high sensitivity and are commonly used to probe the structure and dynamics of large proteins and molecular complexes ^45^. NaK has 32 methyl bearing Ile, Leu, and Val residues (out of 110 total residues) for 53 probes per monomer in total, which are dispersed rather uniformly throughout the structure. The methyl groups were assigned previously ^46^. We then compared the methyl chemical shifts for U-[^15^N^12^C^2^H] Ile (δ1), Leu/Val labelled NaKΔ18 and NaK FL. The isotopic labeling of the methyl groups in these samples is a racemic mixture of ^13^CH_3_/^12^C^2^H_3_ for each Leu Cδ1/δ2 and Val Cγ1/γ2, respectively (referred to as ILV-labeling throughout). The methyl CSPs are generally less than 0.05 ppm, with only a few residues at the proposed ‘hinge region’ ^2,47^ and near the HBC showing larger perturbations (Figure 2).

**Figure 2.**
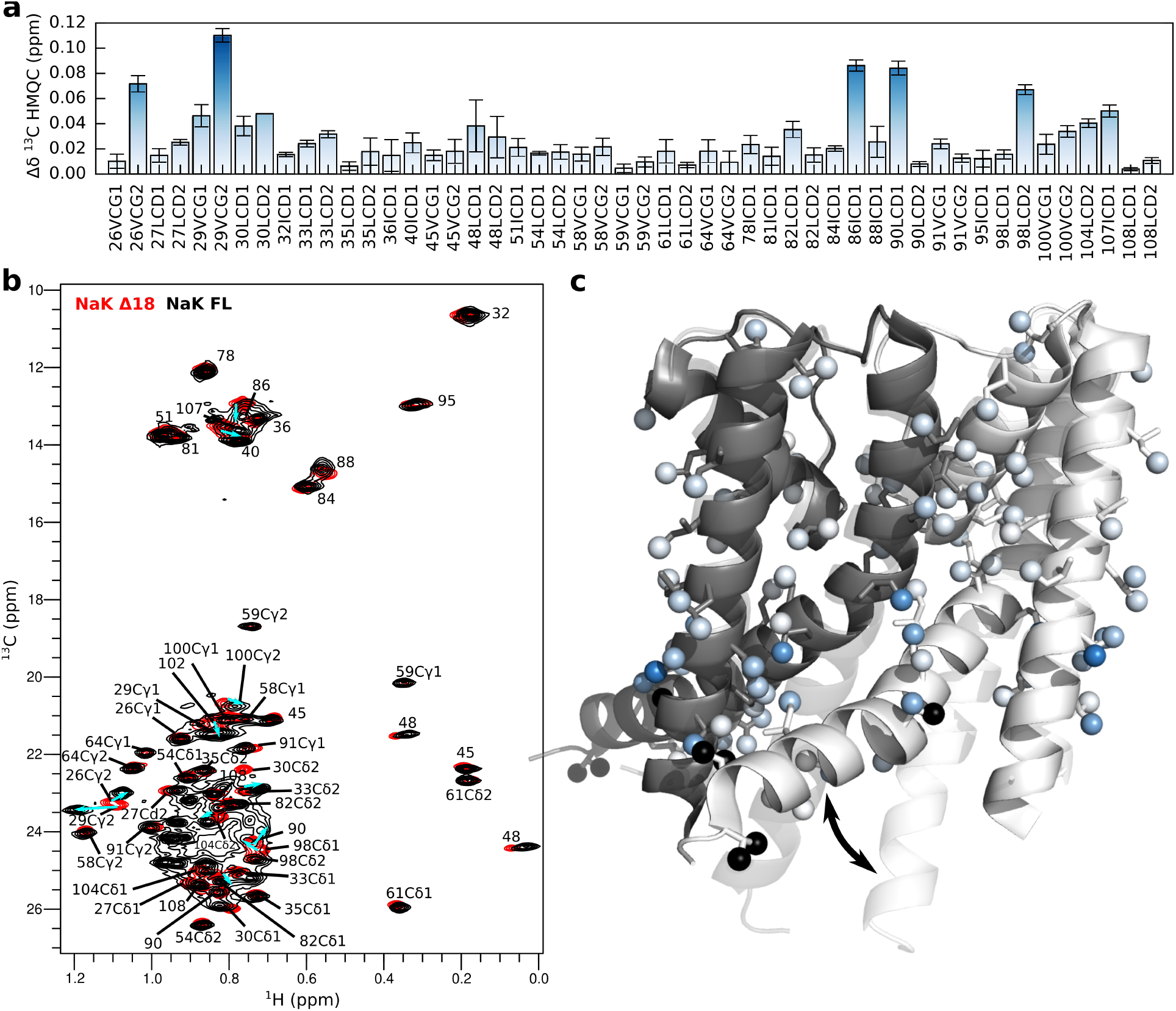
Methyl chemical shift perturbation between NaKΔ18 and NaK FL. a) Combined ^1^H,^13^C chemical shift difference extracted from the ^1^H,^13^C-HMQC spectra shown in (b) for NaKΔ18 (red) and NaK FL (black). Chemical shift assignments are shown in (b) with cyan arrows highlighting more pronounced perturbations. c) Chemical shift differences plotted on the ILV methyl groups of the NaKΔ19 crystal structure (3E8H) in the same color scale as panel (a). Black spheres indicate unassigned residues. NaK FL crystal structure (PDB: 2AHZ) shown in faded overlay, with arrows highlighting the M_2_ helix conformational change between the “open” and “closed” conformations.

In accordance to previously published NaK FL solid state NMR data ^43,46^ we observe broadening of resonances around the hinge, which becomes more severe in NaK FL. Whereas residues in the M_2_ helix below V91 could not be assigned in ssNMR data on NaK FL, all of the M_2_ resonances are observable by solution NMR in bicelles. There is broadening of both the amide and methyl resonances of G89 and L90, along with the L98 methyl resonance (Figure 2, Figure SI2), but resonances downstream of the hinge region, such as the Cγ1/2 methyls of V100 and V102, do not appear to be broadened. As we have previously reported ^46^, the flexibility of the M_2_ helix in the region V102-L108 has increased ps-ns timescale dynamics, which could explain why this region is not observable in the solid-state data. The data also suggest that HBC has enhanced dynamics in different lipid environments (DMPC/DHPC bicelles here and Lewis et al.^46^, asolectin bilayers in Shi et al. ^43^). The ability to observe the terminal parts of M_1_ and M_2_ helices using solution NMR gives us a unique opportunity to probe the structure of the inner gate of a K^+^-like channel more directly by acquiring NMR distance restraints.

### Open NaKΔ18 conformation in lipids, consistent with X-ray structure

Since the chemical shift differences between NaKΔ18 and FL appear rather small, we wanted to more clearly establish the specific structural differences at the HBC between the two variants and whether the observed changes really reflect the proposed transition between closed and open conformations in these constructs. Although chemical shifts directly reflect the shielding of a nucleus by the surrounding electrons and thus report on structure, they have a non-linear dependence on multiple structural features that makes direct interpretation of chemical shift perturbations challenging. For example, methyl CSPs may report on changes in bond angles, polarity, aromatic ring currents, change in populations of dynamically interconverting structures, and other factors. Amide chemical shifts have additional dependence on backbone torsion angles and hydrogen bonding. To more directly compare the structures of the two NaK variants, we decided to measure distances between CH_3_ groups of ILV labeled NaK samples using NOESY NMR experiments. Such distance measurements are one of the most precise and well-established NMR techniques and are routinely used to solve structures by NMR ^48^.

Very high-quality methyl NOESY spectra could be obtained for each of the NaK variants in DMPC:DHPC isotropic bicelles (q=0.33) at 40 °C, displaying between 250-325 off-diagonal cross-peaks in total. A mixing time of 250ms was used, which should lead to observation of the off-diagonal cross-peaks between methyl groups that are within 10Å. We did not observe all of the expected possible inter-methyl cross-peaks in the NaK crystal structure in this distance range, but 110 of these “missing” peaks are expected to be weak (79 of these unobserved cross-peaks report on atoms separated by more that 8Å) or would overlap with the diagonal peaks in our 3D experiments (31 crosspeaks had too small separation of their diagonal ^13^C frequencies). The absence of peaks originating from spin diffusion was further verified by comparing the spectra of ILV labeled NaK recorded in both D_2_O, and H_2_O ^46^, and confirms that the NOESY cross-peaks correspond directly to the distance between methyl groups.

NaKΔ18 crystallizes in an open conformation (PDB 3E8H), and we used this 1.8Å resolution structure as a model for the open HBC conformation when analyzing our NMR data. We used the NaK FL structure (2AHZ, 2.8Å resolution) as the model for NaK with a closed HBC. Three C-terminal residues, S106-L108, which are missing in the 2AHZ structure were modeled using I-TASSER ^49^. The open bundle crossing conformation captured in the NaKΔ18 structure is assumed to be the result of crystal contacts and to not represent a major conformation of the protein ^21^. Thus, we expected that NaKΔ18 in lipid bicelles, as in our NMR samples, would be in a conformation where the HBC is closed.

The NMR-observed distance restraints for NaKΔ18 in DMPC:DHPC isotropic bicelles are shown on the closed HBC structure in Fig. 3d. Clearly, there is a significant discrepancy between this crystal structure and the NMR distance restraints in the HBC region. NOESY cross-peak intensities are strongly distance dependent, resulting in strong cross-peaks if they are separated by less than 5Å and becoming very weak as the distance approaches ~10Å. 63 pairs of methyls for which we observe strong or intermediate NOESY cross-peaks, would be separated by distances longer than 11Å (many of them by ≈20Å) in the closed crystal structure and 13 strong NOESY cross-peaks are observed for pairs of methyl groups that are separated by more than 8Å. A specific example from a residue near the HBC, I88, makes this discrepancy most evident (Figure 3g). One of the strongest off-diagonal cross-peaks we observe is between I88 and I95, which would be separated by 8.5Å in a closed conformation. However, if the HBC was actually in the open conformation captured in the NaKΔ19 crystal structure these methyl groups would be separated by only half that distance (Figure 3c).

**Figure 3.**
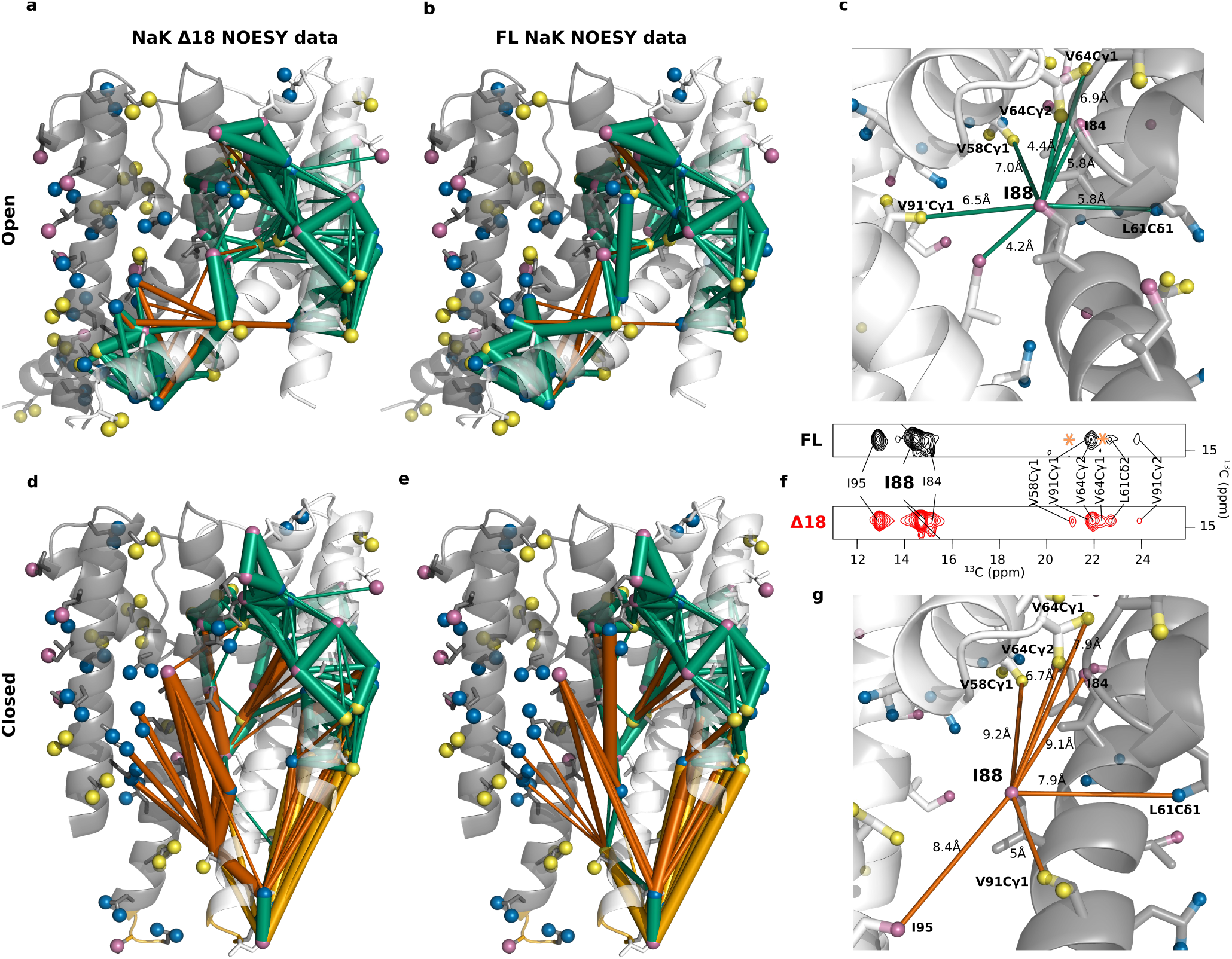
NaK methyl-methyl distance restraints reveal an open HBC in bicelles. (a,d) NaKΔ18 and (b,e) NaK FL methyl-methyl distance restraints measured with using 3D CCH HMQC-NOESY-HMQC NMR experiments on ^13^CH_3_-ILV-labeled protein in q=0.33 DMPC/DHPC isotropic bicelles. The data are plotted on the NaKΔ18 “open” structure (PDB 3E8H) in (a, b) and the same data is plotted on the NaK FL “closed” structure (PDB 2AHZ) in (d, e). Lines connect methyl groups that are close in space based on the NOESY cross-peaks. Line thickness corresponds to the intensity of NOESY cross-peaks. If the methyl groups with an observable NOE are within 10Å in the crystal structure, the lines are colored green indicating consistency between the NOESY data and the crystal structure. If the methyl-methyl distance is >10Å in the crystal structure, the lines are colored orange to indicate inconsistency with the NOESY data. A lighter orange is used to plot NOEs connecting residues missing from the NaK FL crystal structure that were added using I-Tasser. (f) NOESY crosspeaks observed for I88 in NaKΔ18 and NaK FL and their correspondence to (c) open and (g) closed structures illustrate the consistency in the NOE pattern in both constructs, which contrasts with significant distance changes in the crystal structures. Green and red lines indicate distances which are consistent and inconsistent with NOESY cross-peak intensities (as illustrated in f), respectively. Valine CH_3_ groups are colored in yellow, leucines in blue and isoleucines in magenta.

Replotting all of the methyl NOESY data for NaKΔ18 on the open crystal structure shows a striking agreement (Figure 3a). For example, even side chains on the surface of NaK that do not appear to be constrained by packing display strong consistency between the observed NOE data for NaK in bicelles and the crystallographic distances indicating that they are in similar orientations (i.e. V29) (SI Figure 3). The few remaining discrepancies (Figure 3a) can be readily explained by reorientation of the I78 side side chain or a modest differences in the structure or dynamics of the M_2_ helix. This sort of minor discrepancy is not unexpected when comparing protein crystallized from DDM detergent versus protein reconstituted in lipid bicelles.

### Full-length NaK is also “open” in lipid bicelles

The methyl NOESY data establishes that NaKΔ18 exists in an open conformation in lipid bicelles, but what about NaK FL? Although this construct crystallized in a closed HBC conformation, we observe small CSPs between NaKΔ18 and NaK FL in bicelles and need to more directly assess whether these reflect an actual opening-closing transition of the HBC. For this, we have recorded the same 3D ^13^C-NOESY-HMQC experiment on NaK FL. The NOESY cross-peak patterns are very similar between NaKΔ18 and FL, indicating the methyl-methyl distances and structure of the FL channel is nearly identical to NaKΔ18 (Figure 3f, Figure SI4). This is best illustrated by plotting NaK FL NOESY data on the “open” and “closed” structures of NaK (Figure 3b, d). In fact, cross-peaks indicating proximity of N- and C-terminal atoms, a situation which only occurs in the “open” HBC structure, are slightly stronger in NaK FL. Almost all of the same cross-peaks appear in both spectra (with some loss of sensitivity and resolution in NaK FL due to overlap from the additional M_0_ helix residues) and direct comparison of the intensities of each unambiguous NOESY cross-peak between NaKΔ18 and NaK FL (Figure SI5, SI6) shows ratios which are mostly close to 1. This indicates that there are only minor changes in the methyl-methyl distances across the protein. To get an approximate understanding of what distance certain cross-peak intensity changes correspond to, we looked into a well-defined change of the I88 side chain between Na^+^ and K^+^ bound NaK (Figure SI7). There, a 3.5Å change in distance corresponds to ≈3 fold change in NOESY cross-peak intensity, whereas open to close transition would require distance changes of 20Å or more. Although the cross-peaks reporting on HBC structure are strongly consistent with an open FL structure, we do observe difference in the NOESY data for some M_2_ helix residues closer to the selectivity filter. For example, I88, which has been suggested to be important for channel activation in MthK appears to have weaker cross-peak intensities to the selectivity filter and pore helix residues V64 and V58 (Figure 3f, indicated by *) in NaK FL. However, the resolution of our data does not permit more detailed refinement of the structure of M_2_ helix in NaK FL.

In combination, the NOESY and chemical shift perturbation NMR data of NaKΔ18 and NaK FL leave little doubt that, in contrast to X-ray structures solved in DDM, both NaK variants exist in an open conformation in a lipid environment. Since our measurements were performed at 40°C, slightly above physiological temperature, we wanted to see if the HBC changes conformation at lower temperatures. Decreasing the temperature to 20°C did not reveal any chemical shift changes in either NaKΔ18 or FL, that were significant enough to correspond to a conformational change of HBC closing (Figure SI 8).

As we clearly see only a single set of peaks on a HMQC or HSQC spectra, this indicates that unlike in KcsA ^11,12^, only a single major conformation exists in NaK. We have thus tried to detect any minor conformations which could potentially correspond to the closed HBC conformation of NaK. However, we could not detect such evidence in the vicinity of the HBC on the millisecond time scale (a timescale at which conformational changes of such magnitude would be expected) in neither NaKΔ18 ^46^ nor in NaK FL (Figure SI 9). We do observe somewhat enhanced millisecond dynamics for I88 Cδ1 methyl group, which is in line with the broadening we observe for G89 and L90, indicating some conformational dynamics in this region.

### Reconciling NaK structure and function

To assess the functional state of the channel we turned to solid-supported membrane (SSM) electrophysiology ^50,51^. Prior electrophysiology studies of NaK all used the F92A mutant (Figure 1a, b) ^22^, which substantially increases channel flux. SSM electrophysiology was designed to enable study of low-flux channels and transporters and allows us to functionally characterize the NaK channel both with and without the F92A mutation ^51^.For SSME measurements, each NaK variant was reconstituted in 3:1 POPC:POPG liposomes and immobilized on a Nanion sensor. The proteoliposomes had 0.5mM internal K^+^ and 99.5 mM NMDG^+^(N-Methyl-D-glucamine) in 50 mM pH 7 Tris-MOPS buffer. To measure whether the channels are conductive, we then flowed in buffer containing 5mM K^+^, 95 mM NMDG^+^ in pH 7 MOPS-Tris buffer to create a 10-fold potassium gradient. Under these conditions, K^+^ ion transport into the liposomes is expected to trigger positive capacitive currents. Measurements performed on liposomes containing different protein to lipid ratios confirm that we observe currents due to transport and not ion biniding (Figure SI 10). In accordance with the previous electrophysiology measurements ^7,21,22,52^, neither wild-type NaKΔ18 nor FL channels displayed detectable currents, while both of the F92A variants did (Figure 4a). However, the currents observed for F92A-NaK FL were only slightly more than 2-fold smaller than F92A-NaKΔ18, indicating a large population of full-length channels are open and conductive.

**Figure 4.**
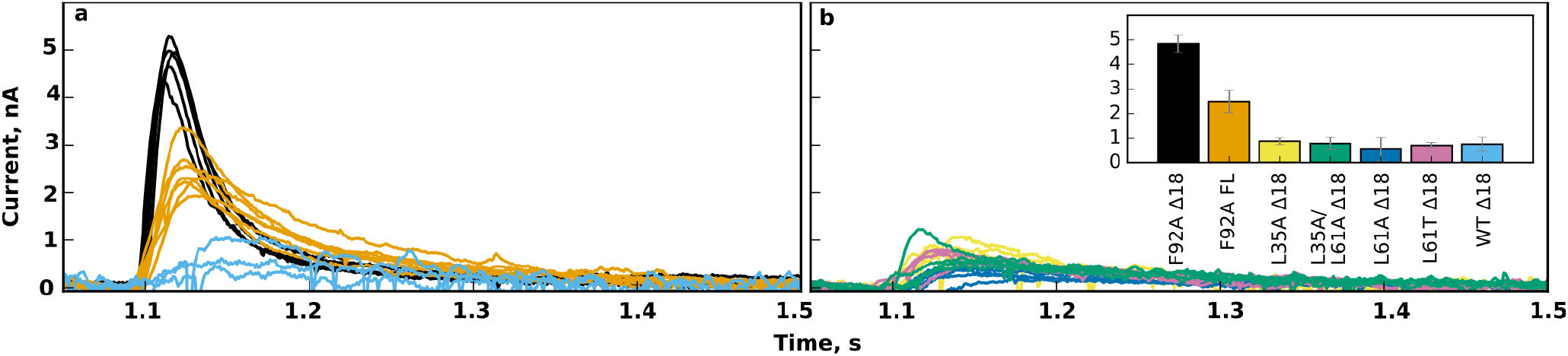
Only F92A mutation induces significant current through NaK. a) Solid-supported membrane electrophysiology recordings of current induced in NaK proteoliposomes by a 10-fold K^+^ gradient. a) Current traces for F92A-NaKΔ18 (black), F92A-NaK FL (orange) and WT NaKΔ18 (blue). Each trace represents the capacitive current from an individual sensor. b) Current traces for NaKΔ18 mutants L35A (yellow), L61A (blue), L35A+L61A (green), L61T (pink). Insert – average peak current and standard error for each construct shown in the same color as the raw current traces.

### How does F92A increase channel flux?

The NMR data indicates that NaKΔ18 and NaK FL have essentially identical open structures of the HBC in a lipid environment, but minor structural differences in the M_2_ helix closer to the selectivity filter. The electrophysiology data show that neither channel is conductive without the F92A mutation, and with this mutation there are relatively modest differences in flux between the FL and truncated channel ^22^. Together, this suggests that the HBC does not perform the main role in regulating NaK flux. We therefore considered whether NaK is gated at the selectivity filter. Indeed, residues in proximity of F92, and not the HBC, have recently been suggested by MD simulations to be the principal players in activation of channels such as MthK and KcsA ^32,35,36,53^, with NaK2K F92A displaying similar behaviour ^35^. In addition, a phenylalanine residue situated just below the P-helix appears to also play a major role in C-type inactivation of KcsA ^23,37^. While a phenylalanine residue in the M_2_ helix just below the selectivity filter is generally conserved in NaK and many K^+^ channels (Figure 1b), F92 in NaK is shifted downwards by one position and is located where a small chain amino acid would be situated in most other channels. The cavity just below the selectivity filter is therefore very narrow in NaK and this was suggested to account for low flux of the wild-type NaK channel ^41^.

As there are no experimental structures of F92A-NaK, we wanted to investigate whether there is allosteric coupling between F92 and the selectivity filter, as proposed for other K^+^ channels, which could provide a more general alternative explanation for gain of flux in the NaK F92A variant. We attempted to test F92A, F92D and F92N variants for this experiment, but polar substitutions significantly reduced protein expression and disrupted tetramer formation (SI Figure 1). The first rather striking observation is a stark difference in NaKΔ18 F92A α-helix stability. NMR studies of a protein of this size in bicelles require deuteration of the protein to reduce relaxation and improve spectral sensitivity and resolution. This is achieved by expressing the protein in D_2_O, then purifying it in H_2_O where it is hoped that most of the amide groups will back exchange to protons so they can be observed in ^1^H/^15^N correlation spectra. This effectively performs an H/D exchange experiment, with the peak intensity in the ^1^H/^15^N TROSY-HSQC spectrum providing a direct readout of the level of back-exchange that occurred during purification. F92A-NaKΔ18 displays much better back-exchange of the pore helix region, which is otherwise very poorly back exchanged in the wild-type NaKΔ18 channel ^46^ (SI Figure 11). This stands in contrast to what was previously observed for NaK2K in *E. coli* polar lipid extract ^54^. The latter study used K^+^ depleted channels, and this suggests an interesting possibility that K^+^ is required for overall stability of NaK. However, when we tried to incubate NaKΔ18 without K^+^ following the procedure from ref 54, the resulting samples had very little NMR signal, suggesting that the protein had aggregated.

Comparison of 2D methyl HMQC spectra of ILV labeled F92A NaKΔ18 and WT NaKΔ18 reveals CSPs that are clustered around the mutation site and the selectivity filter (Figure SI12), similar to what was observed for F103-KcsA by ssNMR ^37^. This suggests that the overall structure is not strongly perturbed upon F92A mutation and both NaK and KcsA respond similarly to mutation of the corresponding aromatic residue. We see rather large methyl CSPs at residues I95 (~0.4ppm) and V91 (~0.2ppm). In addition, smaller but significant methyl CSPs occur throughout the residues in and adjacent to the selectivity filter (V64, V59, I84), pore helix (L61 and L35), and also L48 (Figure SI12). L48 is not located in the selectivity filter, but its sidechain is in close proximity to residue F69, and its methyl group likely reports on the changes in the upper portion of the SF. I88 also displays a moderate CSPs on Cδ1 (similar to I100 in KcsA ^37^). Therefore, CSP data suggests allosteric coupling between the selectivity filter and F92 in NaK. Based on these NMR observables the behavior appears to be similar to KcsA, where the allosteric link between the selectivity filter and the M_2_ residues below it is well established functionally ^23^. As removal of the phenylalanine ring will alter the impact of ring currents in the vicinity of the mutations, we looked for measurable structural changes upon F92A mutation using methyl NOESY experiments.

The difference of diagonal peak intensities between F92A and WT NaK is generally below 10%, with slight changes in V91, L35, L61, I95, I81 and V29. As most of these residues are located in the region displaying poorest H_2_O/D_2_O back exchange, the decreased peak intensities likely reflect increased dynamics (Figure SI13). Overall the off-diagonal NOESY cross-peak patterns, reflecting on the atom distance in space, are nearly identical between NaK F92A and the wild-type protein (Figure SI13, SI14), further reinforcing the conclusion that there are no large global structural changes. However, focusing on the pore region, we observe a cluster of atoms, which display 2 to 4-fold intensity changes between F92A and wild-type protein (Figure SI13). Certain methyl NOESY cross-peaks reporting on the proximity of the M_2_ and pore helices become stronger, suggesting that in the absence of the bulky aromatic F92 sidechain these two helices do pack more closely (Figure 5). However, I88, which was implicated in the gating of several K^+^ channels such as KcsA and MthK and whose side chain was demonstrated to change conformation depending on the nature of ion in the selectivity filter ^46^ is not affected by F92A substitution. Focusing on the selectivity filter, we notice that the S3-S4 sites of the selectivity filter are also altered slightly, as there are small but significant distance increases between the V64 and V59 methyl groups. NaK does not have any methyl containing probes in the upper section of the selectivity filter, but the CSP of L48 Cδ atoms suggest changes in the selectivity filter at this position as well.

**Figure 5.**
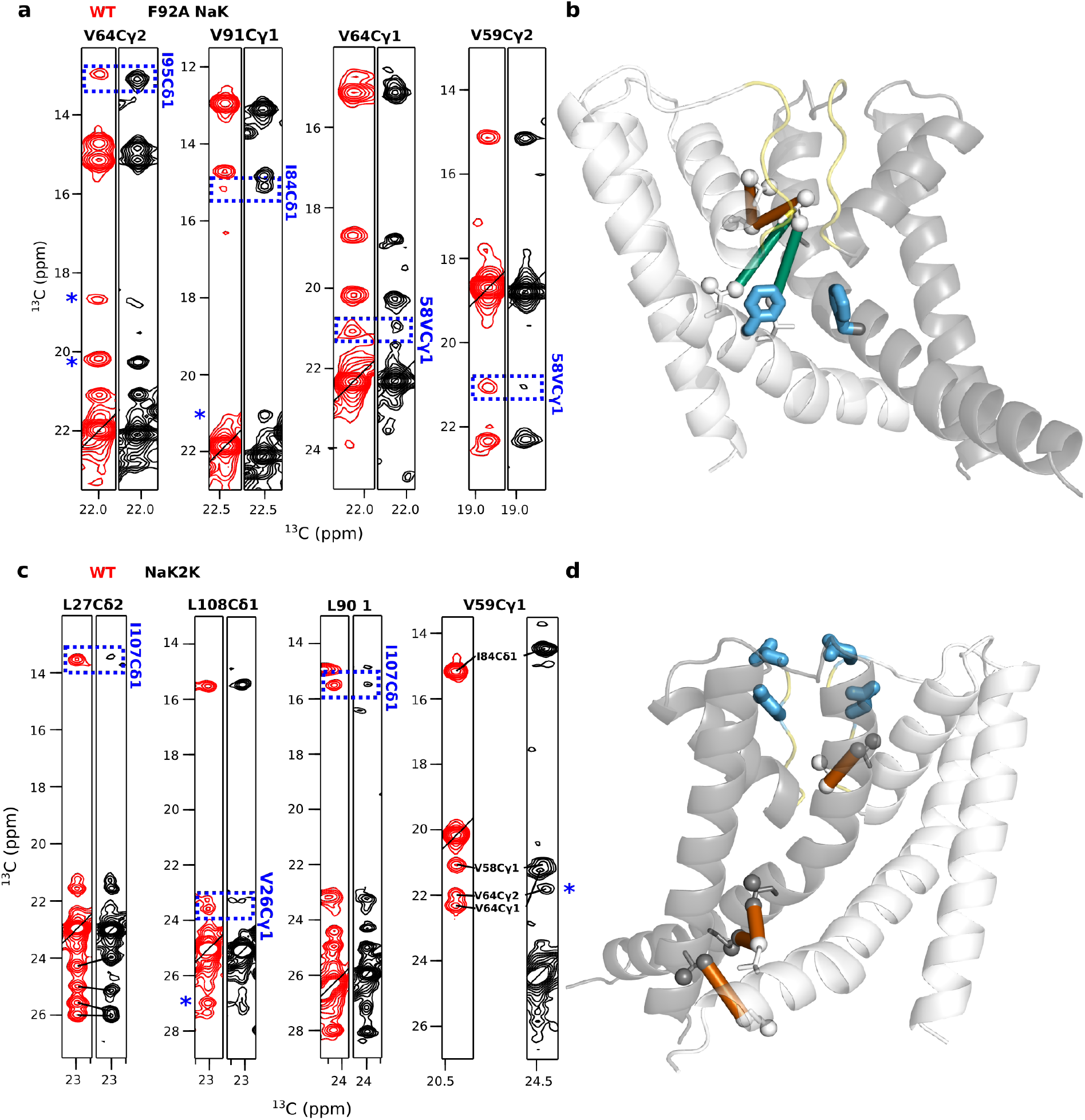
Location of structural changes upon F92A or NaK2K mutation. a) Methyl NOESY cross-peak intensity differences between WT NaKΔ18 (red) and F92A NaKΔ18 (black) are largest (2-3 fold change) for residues located close to the selectivity filer. Blue boxes highlight these changes for NOEs between the methyl group listed in black (top) and blue (side). Blue stars indicated changes which are significant but lower than 2-fold. (b) data from panel (a) is plotted on the crystal structure (PDB 3E8H). Green lines indicate distances which become shorter in F92A NaKΔ18, while orange lines represent distances that become longer. The selectivity filter is colored yellow and mutations are shown in blue sticks. c) Methyl NOESY cross-peak intensity differences between WT NaKΔ18 (red) and NaK2KΔ18 (black). In NaK2K, significant changes are observed in the end of M_2_ helices (blue boxes, labeled as in a), as represented on the structure (d, same color scheme as (b)).

Notably, we do not see much correspondence between the magnitude of chemical shift perturbation and the distance changes between the methyl groups. For example, I95, which displays a very largest CSP of 0.4 ppm does not appear to have exceptional NOESY peak intensity changes with respect to the neighboring methyl-bearing residues. Thus, this large perturbation is likely due simply to ring current changes in F92A. This emphasizes the challenges in interpretation of CSPs.

All of the residues around the selectivity filter displaying significant NOESY intensity changes in F92A-NaKΔ18 (Figure SI13, SI14), especially those located at the interaction point between the M_1_, M_2_ and pore helices (V58, L35, L61, I88, I84, V91), have recently been implicated in channel activation/inactivation and allosteric coupling between the selectivity filter and HBC in MD simulations of KcsA and MthK channels ^32,35,36,53,55^. Moreover, there is some experimental evidence from several different families of K^+^ channels showing that substitutions of these residues can alter ion flux. In KcsA, mutation of L40 (L35 in NaK) to alanine was shown to remove C-type inactivation and activate KcsA at higher pH ^32^; while in the BK channel Slo2.1 a mutation of the corresponding L209T induces constitutive activation ^56,57^. The residue opposite L40 on the pore helix, A73 in KcsA (L61 in NaK) when mutated to E or C leads to a much higher open probability ^33^, while F240T/C substitutions at an identical position in the BK channel Slo2.1 renders the channel maximally active and no longer sensitive to the activator niflumic acid ^56^. The K2P channel TREK-1 responds to a broader range of mutations perturbing the pore helix, where substitutions (G137/V58 (NaK), W275{S/A/T/L/N/Q/D/E}/Y83 (NaK), W276/I81 (NaK), I148/F69 (NaK)) produce channels with increased flux and also uncouple channel opening from stimuli ^58,59^.

We investigated whether altering the structure of pore helix by mutating the aforementioned residues would also activate NaK. To test this, we introduced mutations in NaKΔ18 at positions structurally impacted by F92A in our NMR studies and which were previously shown to increase conductance in KcsA or other channels: L35A, L61A, L35A+L61A and L61T. All of the variants expressed to similar levels and migrated identically on the size-exclusion column as the wild-type NaKΔ18 (Figure SI1) indicating that they retained a tetrameric structure. Nevertheless, increased aggregation propensity suggests that the protein stability is compromised by these mutations. However, unlike F92A-NaKΔ18, none of these mutations resulted in ion fluxes detectable by SSM electrophysiology (Figure 4b).

### Coupling between selectivity filter and HBC

Distance changes between the M_2_ and M_1_ helices and the selectivity filter and pore helix in F92A-NaKΔ18 indicate that the pore helices are coupled to the surrounding structures. Indeed, such a coupling was suggested to play a central role in KcsA C-type inactivation, where the progressive opening of M_2_ helices allosterically transmit the signal causing the selectivity filter of KcsA to collapse, with F103 playing major role in the process ^23,60,61^. In such a mechanism, structural changes of the selectivity filter may also affect residues below the selectivity filter. While it is not clear how general such allosteric coupling mechanism is, NaK2KΔ18 (Figure 1a), a variant of NaK with a 4-site selectivity filter shows widely distributed chemical shift perturbations as observed by NMR ^44^, including the residues comprising M_2_. We therefore investigated if the four-site selectivity filter in NaK2K induces structural changes in the M_2_ helices, for example making the HBC more closed, which are not evident from the X-ray structures.

Our first observation was that NaK2K Δ18 expressed in D_2_O and purified in H_2_O back-exchanges more slowly with water than the wild-type protein, consistent with a more rigid structure (Figure SI15) and in contrast with F92A-NaK. This is further supported by the observation that peak intensities of V58 and V59 become stronger (Figure SI18) in the four-site SF filter containing NaK2K. The broadening of these residues in NaKΔ18 likely occurs due to slow dynamics, and since these residues are not dynamic on the millisecond time scale (Figure SI8), the rigidification most likely happens on the microsecond timescale. Upon close examination of methyl ^1^H/^13^C-HMQC spectra, we noticed dramatic shifts (>1ppm) for residues in and near the selectivity filter (V64, V59) (Figure SI16, SI17) that likely arise simply from ring currents due to the D66Y substitution in NaK2K that is in close proximity to these residues (Figure SI17). We also observe significant changes in the side chains of the residues just below the selectivity filter, including the hinge region (I88, L90, L91, L98) and even more distant residues such as V100, in agreement with previous solution amide (^15^HN) ^44^ data. The shifts are of comparable or larger magnitude than the perturbations we observe in F92A NaKΔ18.

After recording and analyzing methyl NOESY data for NaK2K, we observe that NOESY cross-peak intensities remain largely the same between this variant and NaKΔ18. Out of 241 off-diagonal crosspeaks we assigned, only 11 display changes in intensity larger than two-fold. In spite of the dramatic chemical shift perturbations, we only detect subtle changes in the pore helix, as reported by V59 or L35 residues (Figure in 5c, d). In addition, we also observe some changes in NOESY cross-peak intensities in the termini of the M_2_ helices (Figure 5c, d, Figure SI18, SI19). For example, L108 displays weaker cross-peaks to its neighbors in NaK2K and L27 and L90 shows two-fold weaker cross-peaks to I107. Remaining small but significant changes between M_2_ helix contacts supports the observation of selectivity filter – HBC coupling, although the actual structural changes might be smaller than expected.

## DISCUSSION

Accurate assignment of X-ray and cryoEM structures to functional states is of crucial importance, since these structures form the basis for more detailed mechanistic models, are the starting point for MD simulations, and influence the design of future experiments. The difficulty of this problem can be illustrated by ion channels, where X-ray structures of K^+^ channels from various families were integrated into a single HBC steric gating model as a general mechanism of ion channel activation, yet subsequent investigations demonstrated the HBC does not pose a barrier for K^+^ permeation under conditions where the channel is non-conductive in many cases ^16,20,25,27–33,62^. Structural changes occurring in the HBC during the functional cycle of a channel are further obscured by the tendency of the same channel to crystallize in only one conformation (either open, like MthK, or closed, like KcsA or Kir channels). KcsA is among the best studied channels by X-ray crystallography, yet among more than 100 structures deposited to the Protein Data Bank, there is not a single structure of the open wild-type protein. Other methods, such as EPR or SANS appear to detect structural changes at low pH ^63–65^. As none of the deposited structures are solved below pH of 4.0, the pH at which the HBC in KcsA was suggested to open, it might be that the open conformation of KcsA displays substantial heterogeneity which precludes crystallization. Furthermore, removal of salt-bridges stabilizing the closed state of KcsA by mutagenesis results in a heterogenous set of crystals, each displaying a different degree of HBC opening ^6^. In agreement with such interpretation, NaK solution and solid-state NMR data suggests that the C-terminal half of the M_2_ helix displays enhanced dynamics ^43,46^. Moreover, the lack of X-ray structures of certain intermediates and the consequent difficulty in reconciling these structures with functional data is not limited to KcsA. For example, BK Slo2.2 and K_2_P TASK channel cryoelectron microscopy structures have suggested the gate to be at the HBC ^18,19,66^, while this localization of the gate appears to disagree with functional observations ^20,28^. It also extends away from K+ channels, to channels such as pLGICs ^67^.

We have measured NMR spectra of the full-length NaK and its N-terminally truncated variant, NaKΔ18, which is missing the M_0_ helix. While the X-ray structures show two very different “open” and “closed” conformations for these variants, amide and methyl 2D NMR spectra indicate moderate chemical shift differences between the two protein constructs in lipid bicelles. We show that the most pronounced methyl chemical shift perturbations, which are commonly associated with the largest structural changes, often occur due to changes in the orientation of neighboring aromatic side chains (as can be seen in the 2D ^13^C HMQC spectra of F92A NaK and NaK2K). In this way, large methyl CSPs from aromatics neighboring side chains might “conceal” a more important change in structure with a lower CSP. In other words, the methyl groups chemical shifts over-represent reorientations of the aromatic side chains. To better characterize FL NaK and NaKΔ18 structures we turned to NOESY distance measurements between the methyl groups of Leu, Val and Ile. Surprisingly, they suggested that both NaKΔ18 and full-length variants are in the same, “open” conformation in DMPC/DHPC lipid bicelles. Thus, our data adds to the accumulating evidence that K^+^ flux across several ion channels is not hindered by a steric HBC occlusion, and is likely regulated at the selectivity filter instead.

In light of these observations on multiple different channels, the HBC hypothesis was revised to an allosteric model, where physical separation of the C-termini of M_2_ helices allosterically transmits changes to the selectivity filter, causing opening of the channel. Thus, when the orientation of the M_2_ helix is perturbed by opening at the HBC in MthK or KcsA during MD simulations, a small correlated change between residues S69 (S57), V70 (V58) and L40 (L35) in KcsA ^32^, or (I84, T59) in MthK ^35,53^ leads to a slight expansion of ~1Å at the bottom of the selectivity filter, which leads to flux. Similar residues were suggested by analyzing MD trajectories (I56,V91,F87)^36^. This allosteric model is proposed to extend somewhat generally across multiple K^+^ families. Our data and previously published data ^43,46^ on NaK suggests that the termini of M_2_ helices are somewhat flexible on various time-scales, which may correspond to intrinsic swaying of M_2_ helices. Such motions have not been investigated in other channels, such as KcsA or MthK and may be important in understanding the gating mechanism.

While the coupling between the selectivity filter and M_2_ helix residues below it is well delineated in KcsA, it is interesting to note that we observe similar coupling in NaK. Besides our previous observation that the nature of ion in the selectivity filter impacts the M_2_ helices ^46^, here we also observe chemical shift perturbations and NOESY cross-peak intensity changes in F92A NaK and NaK2K that support allosteric signal propagation between the SF and M_2_ helix. This is consistent with the SF gating model, but definitive interpretation is complicated by several factors. Unlike the majority of channels with a tetrameric K^+^ channel fold, F92A in NaK points towards the conductance pathway and occupies a position where a small side chain, A or G, is typically located in other K+ channels (Figure 1b). Indeed, a previous study on MthK made an A88F mutation to mimic NaK and this greatly reduced the observed flux amplitude (although it actually increased the open probability) ^41^. Mutations in this location appear to affect K^+^ channel gating in general ^20,38–40,42^, but not always in the same manner. In TREK-1 similar substitutions (G171F and A286F) reduce flux ^68^, although the wild-type level of flux of this variant can be recovered by adding activator flufenamic acid. Moreover, replacement of L267 (I88 in NaK) in the conduction pathway of Slo2.1 and Slo2.2 to an even bulkier W actually produces constitutively open channels with high activity ^28^, which appears to conflict with the conclusion obtained by Shi et al ^41^. We investigated whether perturbations of the pore helix at positions known to affect some of K^+^ channel gating (which generally coincide with the residues in contact with all three helices: pore, M_1_ and M_2_) can also activate NaK channel with F92 intact. While we could see an effect on the tetramer stability both on SDS PAGE and gel filtration profiles, the L61 and L35 mutants did not display any flux. However, we do not know if such mutations sufficiently affect the selectivity filter and more well design functional and structural studies might help to dissect the overlapping effects that lead to NaK gating at the selectivity filter.

In light of these observations, what is the functional significance of the closed crystal structure obtained for NaK FL? Similar observations have appeared in the literature in the past, where a constitutively open KcsA variant, as demonstrated by single channel recordings and EPR spectroscopy, crystallized in nearly the same conformation as the closed channel ^69^. Crystal lattice forces might explain why NaK FL appears to be closed in the X-ray structure or this state might represent some minor intermediate with yet unassigned function that was stabilized by the particular crystallization conditions ^70^. In these studies, crystal lattice forces were indicated as preventing the channel from opening in the crystal. Similar phenomenon might explain why NaK FL appears to be closed in the X-ray structure. Alternatively, a closed FL NaK structure might represent some minor intermediate with yet unassigned function, which was stabilized by the particular crystallization conditions ^71^. This raises a question: how reliably can K^+^ channel X-ray structures reveal the full repertoire of functionally relevant states? This problem extends beyond just the conformation of the HBC in closed channels and involves the structural heterogeneity of the selectivity filter as well. It was recently demonstrated that the S0-S2 site region of the selectivity filter of NaK displays a high degree of flexibility by NMR ^46^, although it appears to be ordered in the X-ray structures ^7,21^, and only minuscule differences in side-chain orientation are captured by cryoEM ^72^. Similarly X-ray crystallography displays no differences between K^+^ selective NaK2K SF structures under different ionic conditions ^22^, while differences were detected by NMR ^73^. In KcsA, variants with very different modal gating properties displayed identical SF filter conformations crystallographically ^74^, yet differences between the spectra of these variants were detected with NMR ^75^. It is thus clear that although crystal structures have provided invaluable insight into the structure of ion channels, they have revealed only a limited subset of the full range of functional states and better understanding ion channel function will require structural techniques that can probe channel structure in lipid environments and identify the multiple conformations necessary for the dynamic gating and inactivation of these channels.

## METHODS

### Protein expression, purification, and reconstitution

All NaK variants used for this study were cloned into a pET15b vector as described previously ^44^. The sequence of NaKΔ18 construct, including T7 promoter and terminator was:

aaaaaacccctcaagacccgtttagaggccccaaggggttatgctagttattgctcagcggtggcagcagccaactcagcttcctttcgggctttgttaTTCTTTTTTACGATTCGATAATATACTTGGCAATTGTACATTAACTGCTAACTTATGAATAAATCCAAACACTAGTCCAATCCCAATAAATATGTATAAAATTGTAAATATCTTTCCGAAATCAGTTTGCGGACTAAAATTCCCATCACCGACAGTCGTCAACGTGACCACACTAAAATATAAAGCGTCAATAGGACGTAATCCTTCAACTGTACTATAAAAAATCGTACCCGATATTAAAGTCAAAATTGTTAATACAAATAATACTTGAAATTCTTTATCTTTCCAcatatggctgccgcgcggcaccaggccgctgctgtgatgatgatgatgatggctgctgcccatggtatatctccttcttaaagttaaacaaaattatttctagaggggaattgttatccgctcacaattcccctatagtgagtc gtatta

All mutations, except NaK2K were introduced into this plasmid.

NaK FL sequence used:

aaaaaacccctcaagacccgtttagaggccccaaggggttatgctagttattgctcagcggtggcagcagccaactcagcttcctttcgggctttgtta ATTCGATAATATACTTGGCAATTGTACATTAACTGCTAACTTATGAATAAATCCAAACACTAG TCCAATCCCAATAAATATGTATAAAATTGTAAATATCTTTCCGAAATCAGTTTGCGGACTAAA ATTCCCATCACCGACAGTCGTCAACGTGACCACACTAAAATATAAAGCGTCAATAGGACGTA ATCCTTCAACTGTACTATAAAAAATCGTACCCGATATTAAAGTCAAAATTGTTAATACAAATA ATACTTGAAATTCTTTATCTTTCCACGCTCGTAAACAGGCTCTTAACATTCGCTTTAAAGTTA ACAAAAATGAAAGGGCcatatggctgccgcgcggcaccaggccgctgctgtgatgatgatgatgatggctgctgcccatggtatatct ccttcttaaagttaaacaaaattatttctagaggggaattgttatccgctcacaattcccctatagtgagtcgtatta

NaK2K Δ18 sequence:

aaaaaacccctcaagacccgtttagaggccccaaggggttatgctagttattgctcagcggtggcagcagccaactcagcttcctttcgggctttgtta ATTCGATAATATACTTGGCAATTGTACATTAACTGCTAACTTATGAATAAATCCAAACACTAG TCCAATCCCAATAAATATGTATAAAATTGTAAATATCTTTCCGAAATCAGTTTGCGGACTAAA ATCTCCATAACCGACAGTCGTCAACGTGACCACACTAAAATATAAAGCGTCAATAGGACGTA ATCCTTCAACTGTACTATAAAAAATCGTACCCGATATTAAAGTCAAAATTGTTAATACAAATA ATACTTGAAATTCTTTATCTTTCCAcatatggctgccgcgcggcaccaggccgctgctgtgatgatgatgatgatggctgctgc ccatggtatatctccttcttaaagttaaacaaaattatttctagaggggaattgttatccgctcacaattcccctatagtgagtcgtatta

Mutations were introduced into aforementioned plasmids with a modified QuikChange method using Phusion polymerase ^76^. The following primers, synthesized by Integrated DNA Technologies, were used:

F92A: GAATAAATCCcgcCACTAGTCCAATCCCAAT,GGACTAGTGgcgGGATTTATTCATAAGTTAG;
F92D: GAATAAATCCatcCACTAGTCCAATCCCAAT,GGACTAGTGgatGGATTTATTCATAAGTTAG; F92N: GAATAAATCCattCACTAGTCCAATCCCAAT, GGACTAGTGaatGGATTTATTCATAAGTTAG; L61A: GACAGTCGTagcCGTGACCACACTAAAATA, TGGTCACGgctACGACTGTCGGTGATGG; L35A: ACCCGATATagcAGTCAAAATTGTTAATAC, CAATTTTGACTgctATATCGGGTACGATTTTTT; L61T: GACAGTCGTtgtCGTGACCACACTAAAATA, TGGTCACGacaACGACTGTCGGTGATGG

Resulting plasmids were confirmed by sequencing using Functional Biosciences service.

Protein with natural isotopic composition was expressed as described before (Brettman2015). For the expression of isotopically labelled protein a plate containing fresh NaK BL21 (DE3) transformants was washed with LB media and a 5ml preculture innoculated with an OD_600_ of 0.3 with 100μg/l of carbenicillin. After 1h the resulting LB culture with an OD_600_ of 0.9-1 was transferred to 30ml M9 media supplemented with BME vitamins (Sigma-Aldrich), 5μg/l thiamine (Research Products International) and 1g/l ^15^NH_4_Cl (Cambridge Isotope Laboratories) and 100μg/l of carbenicillin. When the OD_600_ reached 0.9, the cells were pelleted and resuspended in 60ml M9 media in D_2_O (Sigma-Aldrich) containing ^12^C, ^2^H glucose (Cambridge Isotope Laboratories) at 4g/l concentration and 0.25-0.5 g/l ^12^C, ^2^H, ^15^N Isogro (Sigma-Aldrich) and 100μg/l carbenicillin. After this preculture reached OD_600_ of 0.9 it was transferred to 240ml M9 D_2_O containing all the constituents as above in 3L baffled flask. This small modification of the original protocol (Bretman2015) allowed us to obtain 3-fold higher yield of protein from the same expression volume. All the growth steps were performed at 37C with 200-250rpm shaking.

After the culture reached OD_600_ of 0.8, 140mg/l of 2-keto-3-(methyl-d3)-butyric acid-4-^13^C, 3-d, and ~70 mg/l of 2-ketobutyric acid-4-^13^C,3,3-d_2_ were added and incubated for additional 0.5-1 hours. Cold shock (4° C ice slurry) was performed for 10 minutes prior to addition of 0.4mM IPTG. The culture was incubated overnight at 25° C.

Purification and reconstitution of the protein in d54-DMPC and d22-DHPC (Avanti Polar Lipids) q=0.33 bicelles was performed as exactly described previously ^46^.

Briefly, the cells were sonicated in 20x pellet mass (w/v) ml of 100mM NaCl, 200mM KCl, 2.5mM MgSO_4_, 20mM Tris-NaOH pH 7.5, DNase, 1μg/ml pepstatin, 10μM leupeptin, 100μM PMSF, 250mM sucrose and 1mg/ml lysozyme, the insoluble pellet was collected after centrifuging at 30,000g in JA-20 rotor (Beckman Coulter) for 1h, at 4° C. The pellet was ground with homogenizer in 10x (w/v) mass of 100mM NaCl, 200mM KCl 2.5mM MgSO_4_, 20mM Tris-NaOH pH 7.5, 20mM DM and agitated for 4-15 hours at room temperature. Insoluble residue was removed by centrifugation at 30,000g for 15 minutes at room temperature. Soluble fraction was purified using Talon affinity resin (Takara). The resin was equilibrated with buffer A (20mM Tris-HCl pH 8, 100mM NaCl, 200mM KCl, 5mM DM) and solubilized protein containing lysate was incubated with the resin for 20 minutes at RT. Afterwards, the resin was washed 20 resin volumes with Buffer A, 10 resin volumes with buffer A + 15mM imidazole and eluted with Buffer A + 500mM imidazole. Elution was concentrated with centricons (Millipore or Sartorius Biolabs) as per manufacturer instructions and loaded on Superdex 200 Increase 10/300 GL size exclusion chromatography columns and purified with a flow rate of 0.5 ml/min. Eluted fractions containing NaK were reconstituted for NMR or solid supported membrane measurements.

### Solid supported membrane electrophysiology

NaK samples for solid supported membrane electrophysiology (SSME) measurements were reconstituted into 3:1 POPC:POPG liposomes largely as described before ^46^. 25mg/ml POPC and POPG (Avanti Polar Lipids) chloroform solutions were mixed with a ratio of 3:1, respectively, and chloroform was evaporated under nitrogen or argon (Airgas) stream. The lipid film was further washed three time with n-pentane (Sigma-Aldrich) and lyophilized overnight. Before reconstitution lipid film was hydrated in NMR buffer (100mM MOPS-NaOH pH 7, 100mM KCl) for 1 hour at room temperatue. 40-70μM protein in DM purified with FPLC was mixed with lipid to achieve 1:200 – 1:50 monomer:lipid molar ratio and incubated for 30 minutes. The first aliquot of Amberllite XAD-2 (Sigma-Aldrich) was then added and sample was further incubated for 1 hour. After addition of the second aliquot the sample was incubated overnight. The last aliquot was added in the next morning and sample was incubated for 1 more hour. All the incubation steps were done at room temperature. Biobeads were strained and the resulting sample was extruded through a 200nm membrane (Avanti Polar Lipids extruder, Whatman Polycarbonate membranes). Nanion measurements were performed immediately after sample preparation. All the proteoliposomes contained similar amount of protein, as confirmed by SDS-PAGE.

All electrophysiology measurements were acquired on Surfe2r N1 instrument (Nanion Technologies GmbH). 5mm Nanion sensors were prepared as recommended by the manufacturer. NaK proteoliposomes were loaded directly from the reconstitution mixture, without diluting them. We have only used sensors which displayed capacitance values of <30 nF and conductance values of ~1.5 nS, as otherwise the results looked inconsistent for some of the variants, such as L35A.

The buffers were prepared in the following way: 100mM MOPS buffer was prepared first and adjusted to pH 7.0 using a 3M Tris base solution. We then mixed this solution 1:1 (v/v) with 200mM NMDG-Cl or KCl. The final solutions for SSME measurements were prepared by mixing the 50mM MOPS-Tris, 100mM XCl (X=NMDG, K) solutions together to obtain buffers A (5mM KCl, 95mM NMDG, 50mM MOPS-Tris pH 7) and B (0.5mM KCl, 99.5mM NMDG, 50mM MOPS-Tris pH 7).

The sensor was washed for 1s with buffer B, then buffer A, then buffer B in sequential fashion. At each step the buffer perfusion rate was 200 μl/s. 3 recordings were obtained for each sensor, or until the traces would overlap completely.

### NMR measurements

0.7-0.9mM ^15^N, ^2^H, ^13^CH_3_-ILV NaK in 100mM MOPS-NaOH pH 7, 100mM KCl, 7% D_2_O, ~50mM d-54 DMPC:~150mM d22-DHPC (q~0.33), 0.5mM DSS samples were measured at 40° C in 5mM Shigemi (Shigemi) tubes. All 3D CCH HMQC-NOESY-HMQC spectra ^77^ were collected on 900 MHz Bruker Avance III spectrometer equipped with cryogenic probes and controlled by Topspin 3.5p7 software. 250ms NOESY mixing time and 1s recycling delay were employed. The number of complex points acquired for all experiments were 1024 (71ms) x 50 (11ms) x90 (20ms). NaKΔ18 and NaK FL were acquired with 16 scans using uniform sampling, while NaK2KΔ18 and F92A NaKΔ18 were acquired using 32 scans with 40% non-uniform Poisson-gap T_2_-weighted sampling.

CPMG experiments on NaKΔ18 and NaK FL were recorded on 800MHz Varian spectrometer controlled by vnmrj 4.2 software with the same sample conditions as described above. ^13^C MQ experiments employed pulse program by Kay and colleagues ^78^. For both experiments T_relax_ was 20ms. For NaK FL, ν_CPMG_ were:

100, 200 (x2), 300, 400, 500, 600, 700, 900, 1000 Hz. For NaKΔ18 the frequencies were: 50, 100, 200 (x2), 300, 450, 600, 800, 900 Hz.

All NMR data was processed using nmrPipe 9.0 ^79^ and visualized with ccpNMR 2.4.2 ^80^. Non-uniformly sampled spectra were processed with SMILE 1.1 ^81^. Noise for NOESY error analysis was extracted with NMRView J 9.2.0 (One Moon Scientific). Assignment of methyl groups was detailed previously ^46^

Chemical shift perturbations from ^13^C SOFAST HMQC ^77^ and ^15^N TROSY HSQC ^82^ spectra were calculated as ΔδCombined=(ΔδH^2^+W_N/C_^2*^Δδ_N/C_^2^)^0.5^, where W_N_=0.101, W_C_=0.251.

Error bars for chemical shift perturbations in Figure 2a were calculated by extracting standard deviations of each cross-peak from ^13^C HMQC recorded on three individual NaKΔ18 samples and two NaK FL samples, and propagated using standard error propagation. As this demonstrates that errors on these spectra are very small, we do not provide error bars for F92A NaK (Figure SI11 a) and NaK2K (Figure SI16 a) since they would not be visible.

Chemical shift perturbation data was analyzed using Python 2.7.11 scripts and plotted with matplotlib package ^83^. 3D structures were generated using molecular visualization package Pymol 1.8.6.0 (Schrodinger, LLC).

Off-diagonal NOESY cross-peaks were assigned manually and peak intensities extracted using ccpNMR software. Peak intensity comparison was performed using in-house Python scripts and the errors propagated using standard error propagation.

Multiple sequence alignment in Figure 1b was generated using Clustal Omega with default presets ^84^ and visualized with Jalview 2.11.

## ACKNOWLEDGEMENTS

We would like to thank Nathan Thomas for suggestions on performing SSME measurements. Research reported in this publication was supported by NIGMS of the National Institutes of Health under award numbers R01GM116047 and R35GM141748. The content is solely the responsibility of the authors and does not necessarily represent the official views of the National Institutes of Health. This study made use of the National Magnetic Resonance Facility at Madison, which is supported by NIH grant R24GM141526.

## AUTHOR CONTRIBUTIONS

V.K. prepared NaK constructs and samples, collected and analyzed NMR and SSME data, wrote the original manuscript, and revised the manuscript. M.T. assisted in NMR data collection and revised the manuscript. K.H.W. supervised the project, acquired funding, and revised the manuscript.

