## Supplementary Figures for "Reassessing the helix bundle crossing model for gating in a non-selective ion channel"

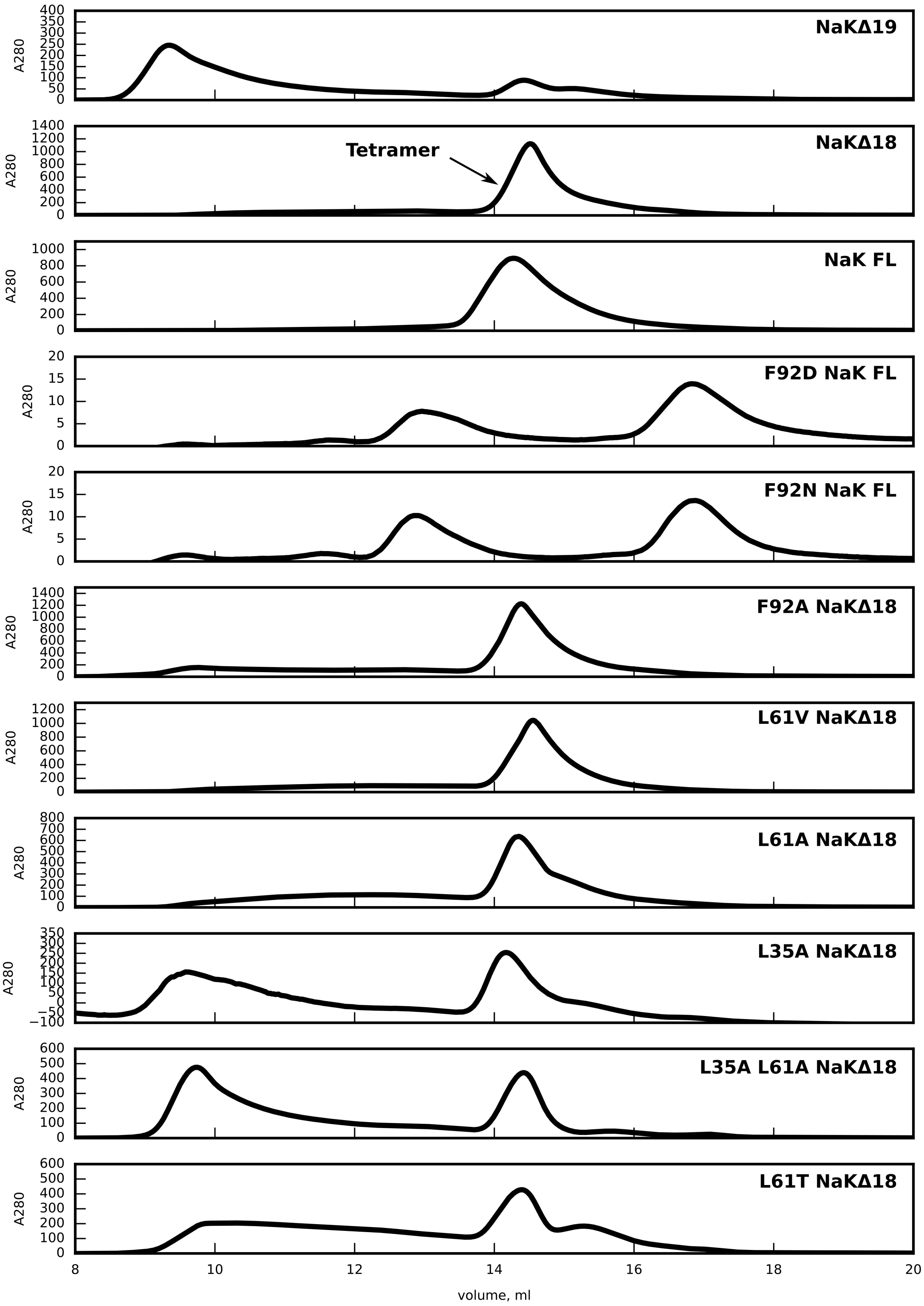


Figure SI1. Size exclusion chromatography profiles of variants used in this study. Top two panels: NaKΔ18 and NaKΔ19 incubated in n-Decyl-β-D-Maltopyranoside for 72h incubation at 4C. The other traces represent samples for which gel filtration was done right after elution from NiNTA. Superdex 200 Increase 10/300 GL columns were used to analyse the samples.

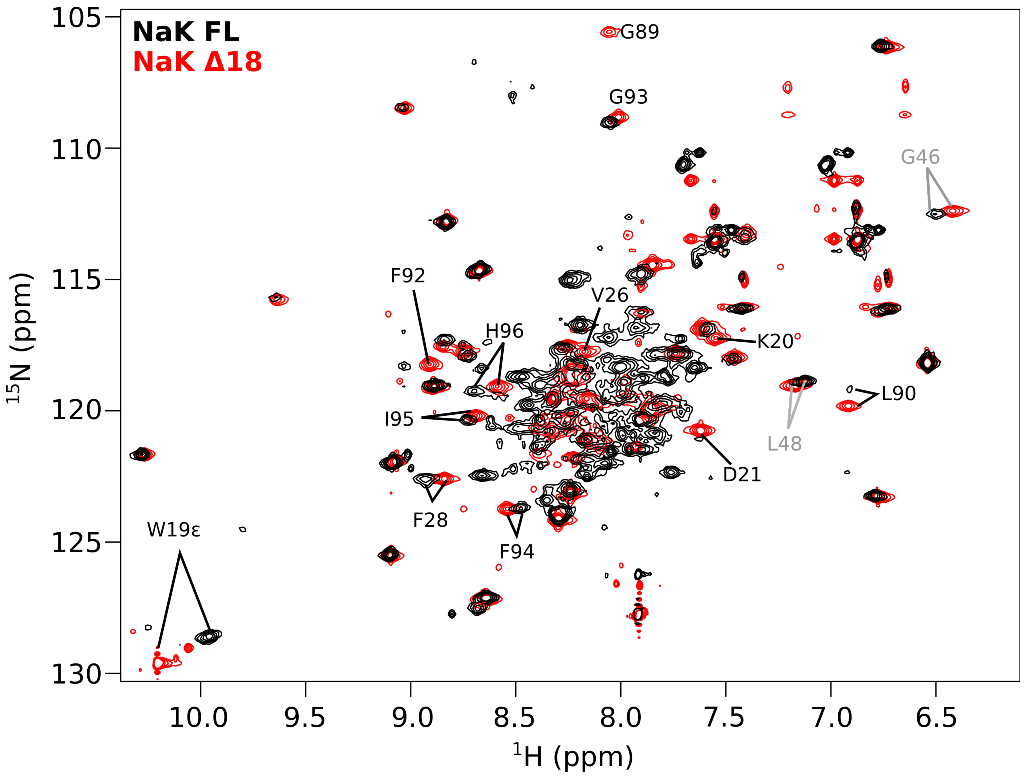


Figure SI 2. Overlay of NaKΔ18 and NaK FL $^{1}$H,$^{15}$N-HSQC spectra. Assignments of non-overlapping residues with more pronounced chemical shift differences are shown.

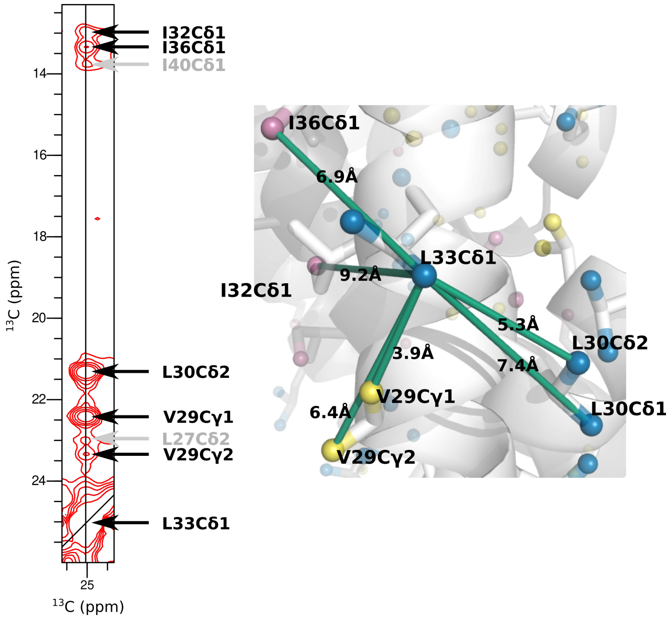


Figure SI3. Correspondence between NMR NOESY and X-ray crystallography data for L33Cδ1. A slice from the $^{13}CH_3$-$^{13}CH_3$ NOESY spectrum of NaKΔ18 showing all the off-diagonal cross-peaks for L33Cδ1, with each cross-peak labeled according to the corresponding atom (left) and the “open” NaKΔ18 structure (PDB 3E8H) centered on L33Cδ1 atom. All the residues within 10Å are connected by lines, and the distance in the X-ray structure displayed on the line (in Ångstroms).

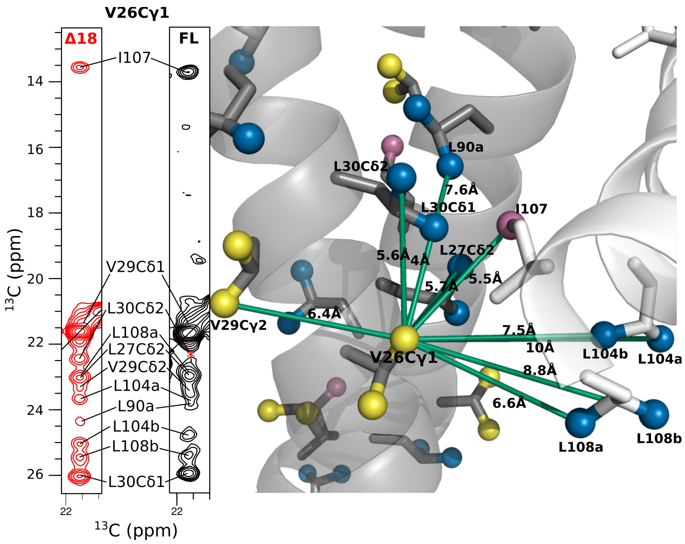


Figure SI4. Correspondence between NMR NOESY data for NaKΔ18, NaK FL and the crystal structure of “open” NaKΔ18 (PDB 3E8H). Slices from the $^{13}CH_3$-$^{13}CH_3$ NOESY spectra of NaKΔ18 (red) and NaK FL (black) with off-diagonal cross-peaks assigned. The data is plotted on the NaKΔ18 “open” structure (right) with all methyl atoms within 10Å connected by lines, and the crystallographic distance shown on the line (in Ångstroms). The red star on NaK FL spectrum denotes the position where L30Cδ2 should appear (this residues is broadened in NaK FL spectra).

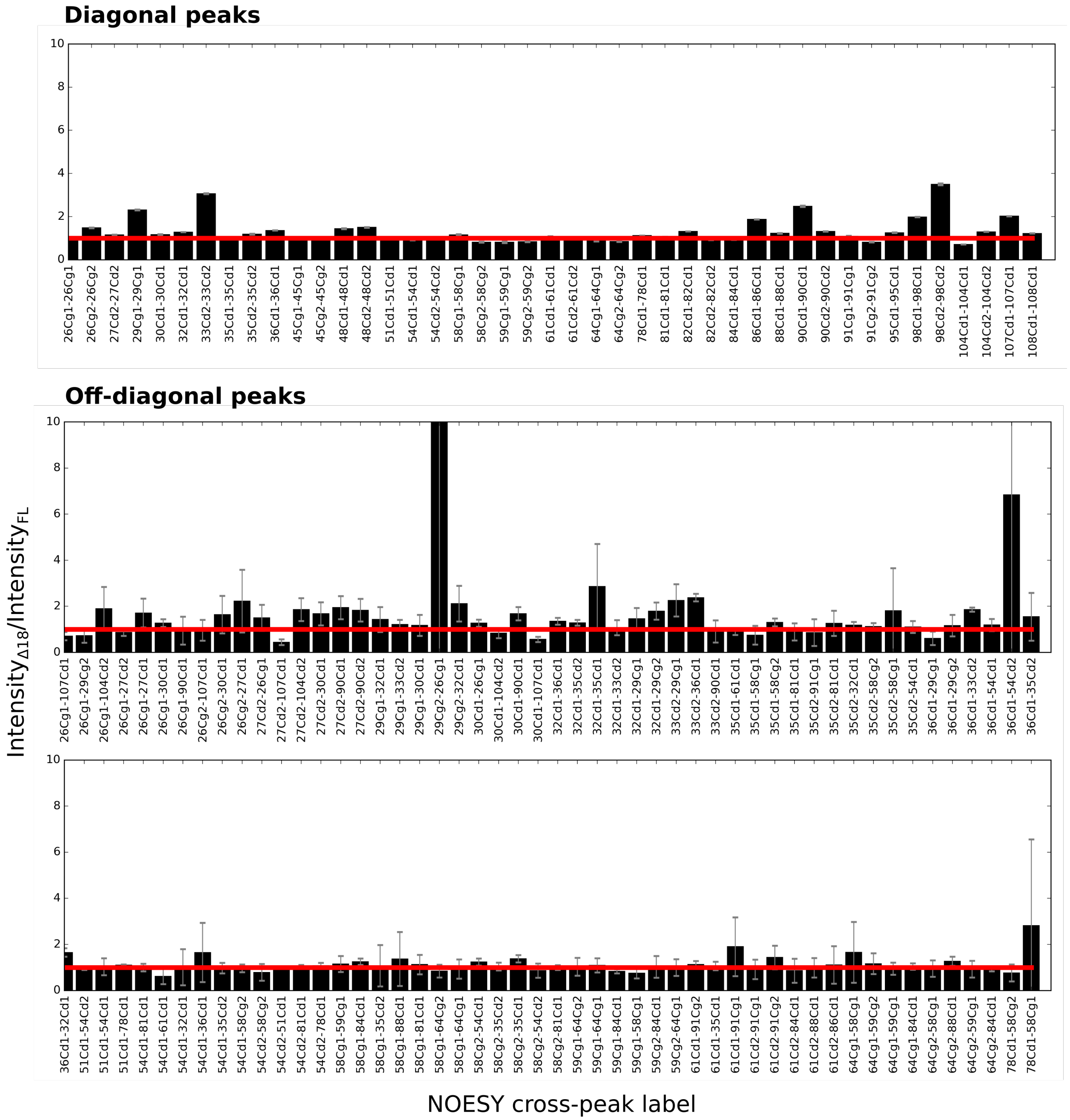


Figure SI5. NaKΔ18/NaK FL intensity ratios of all unambiguous off-diagonal and diagonal cross-peaks plotted with error bars based on the propagated standard deviation of spectral noise. Cross-peak identity is displayed below each bars (name of resonance coming from diagonal peak first). Continued on the next page.

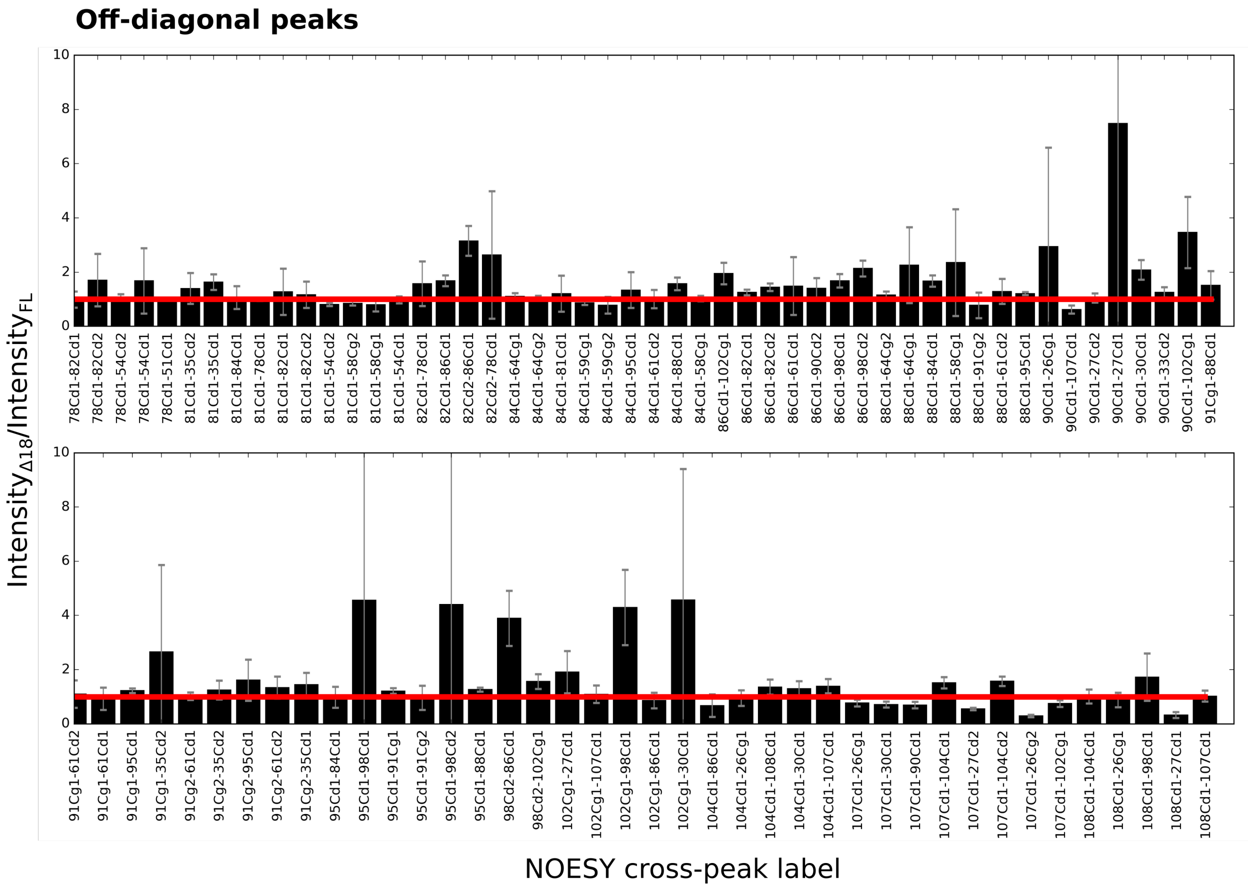


Figure SI5. Continued.

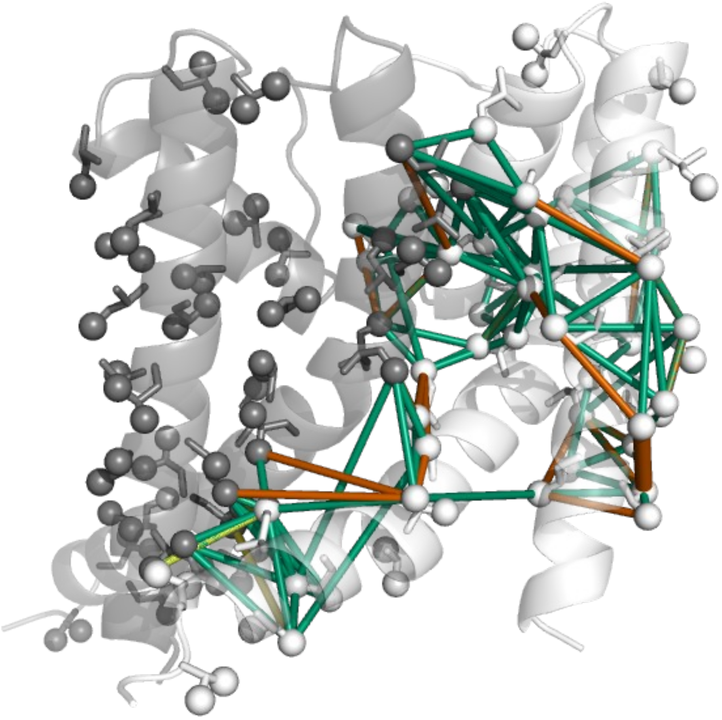

Figure SI6. The same data as in Figure SI5 is plotted on the NaKΔ18 “open” structure. Green lines represent ratios between 0.5 and 2. Orange lines indicate when the peak is more than 2x more intense in NaKΔ18, and yellow lines where the peaks is less than 2x intense in NaKΔ18 as compared to NaK FL.

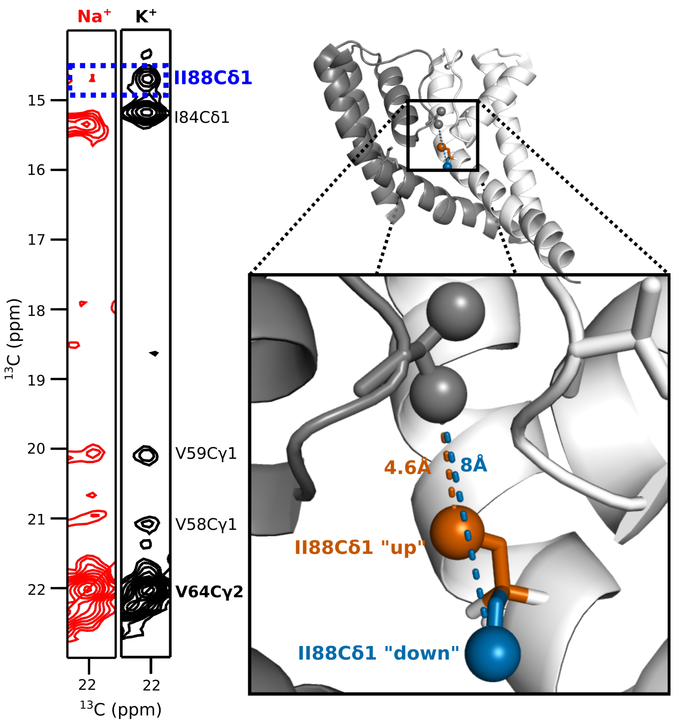


Figure SI7. Impact of ion identity on I88 conformation in NaK. (A) Slices from $^{13}$C NOESY spectra illustrating V64Cγ2 cross peaks in either 100mM (blue) $K^+$ or 600 mM $Na^+$ (brown). The peak with a most pronounced difference between the spectra to I88Cδ1 atom indicated by a dashed box. (B) X-ray structure of NaK (3E8H) with two orientations of I88 and the distance between V64Cγ2 and I88Cδ1 atoms indicated by a dashed line.

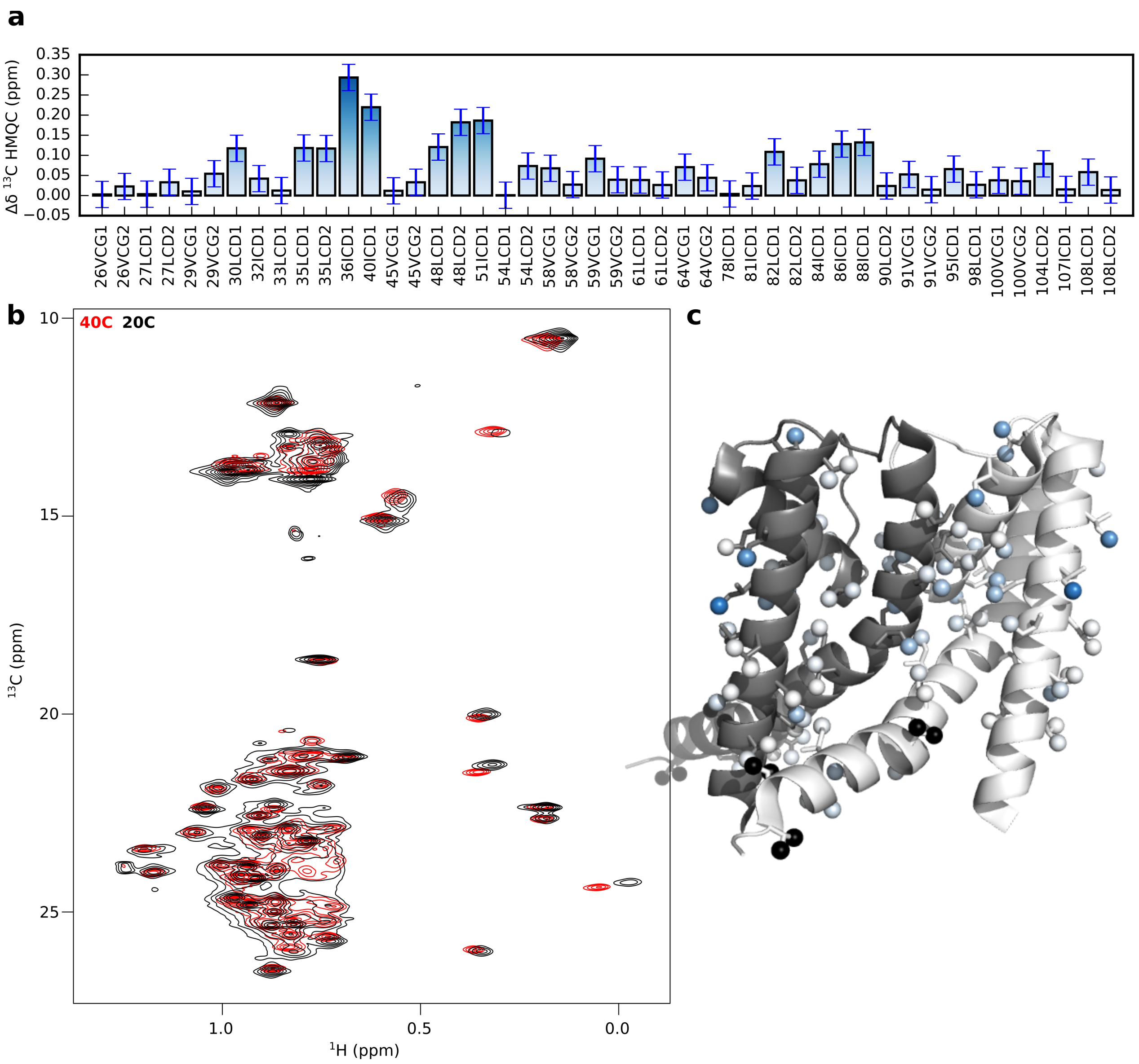


Figure SI8. NaK FL $^{1}H$,$^{13}C$-HMQC spectra at 40°C (red) and 20°C (black). The combined $^{1}H$,$^{13}C$ chemical shift difference is shown as a bar plot (top) and on the methyl groups of NaKΔ19 "open" X-ray structure (right) using the same white to blue color scale. Black spheres indicate unassigned residues. Error bars are based on spectral resolution.

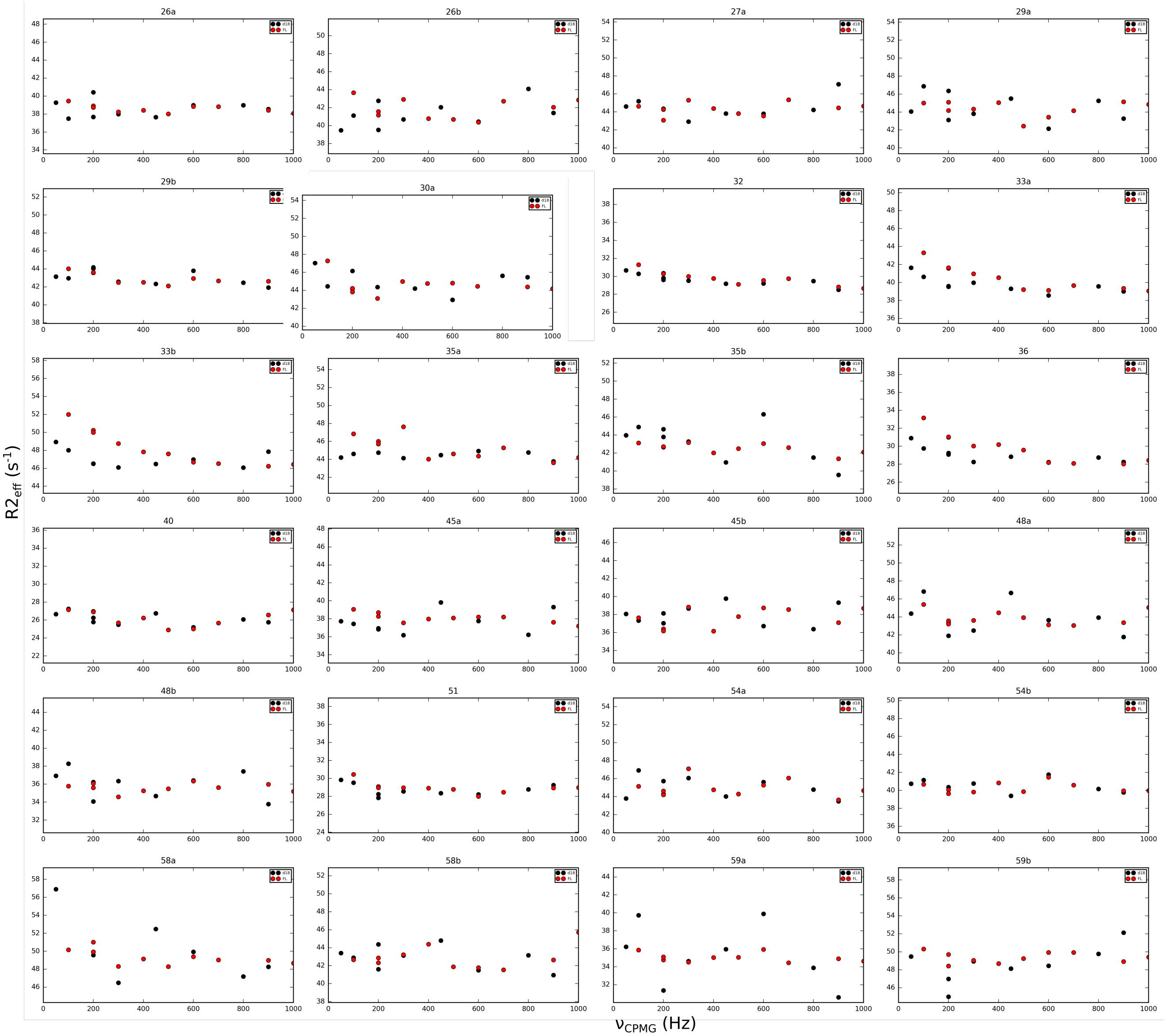


Figure SI9 (continued on the next page). CPMG relaxation dispersion profiles for NaKΔ18 and NaK FL at 800MHz field.

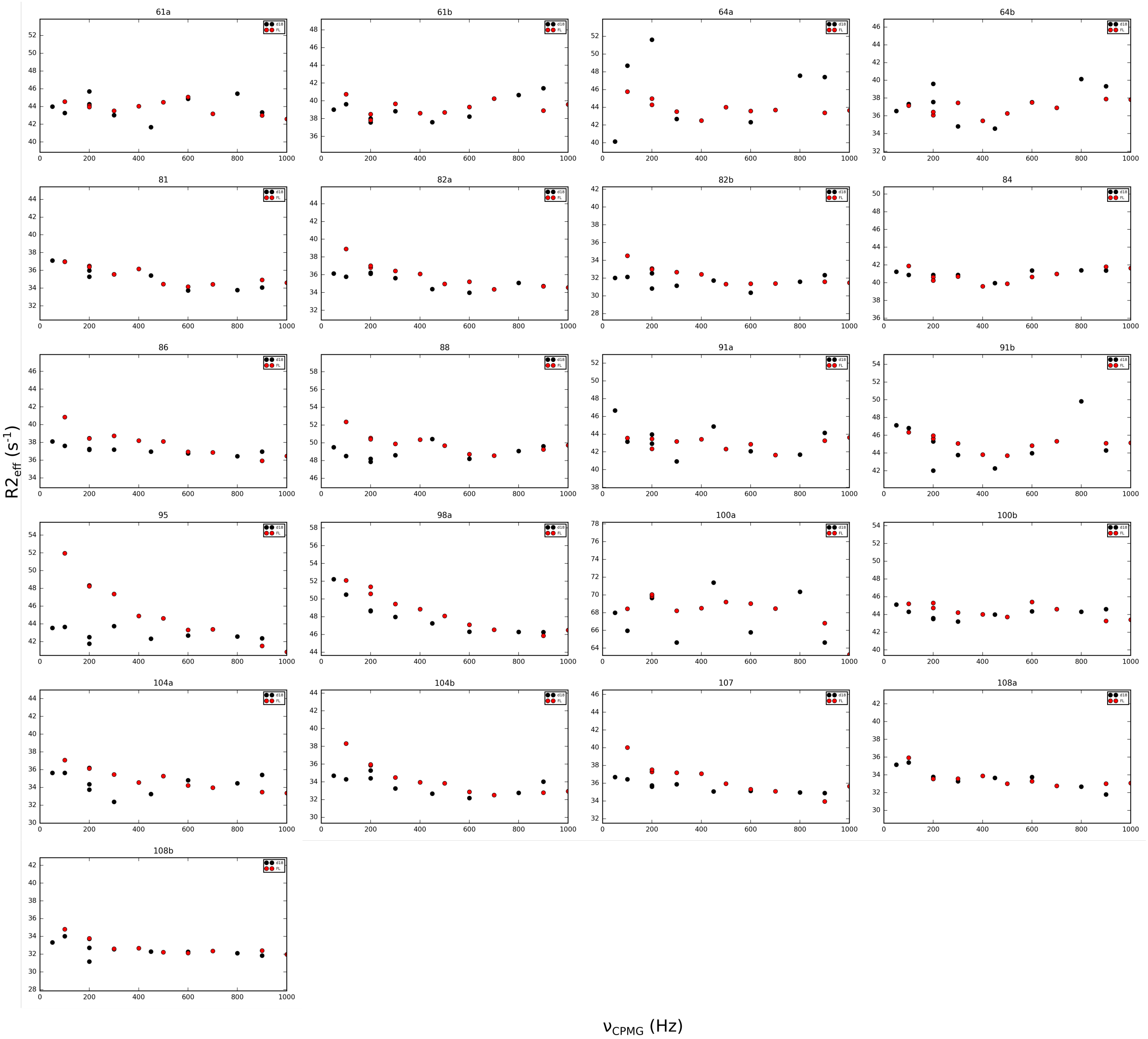


Figure SI9. Continued.

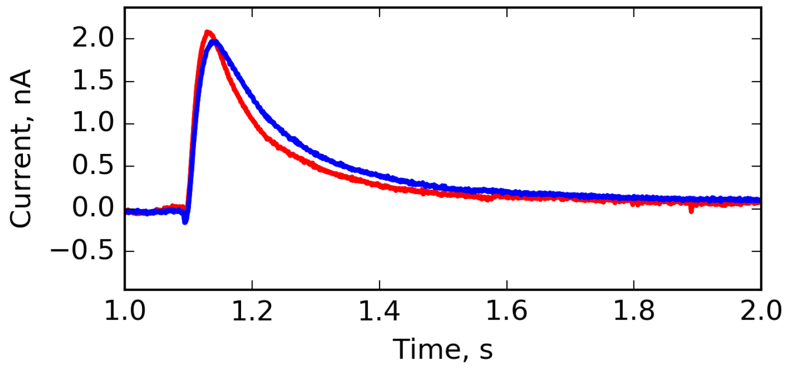


Figure SI10. Comparison of SSME capacitive currents between FL NaK proteoliposomes reconstituted at 1:200 (blue) and 1:800 (red) protein:lipid ratio.

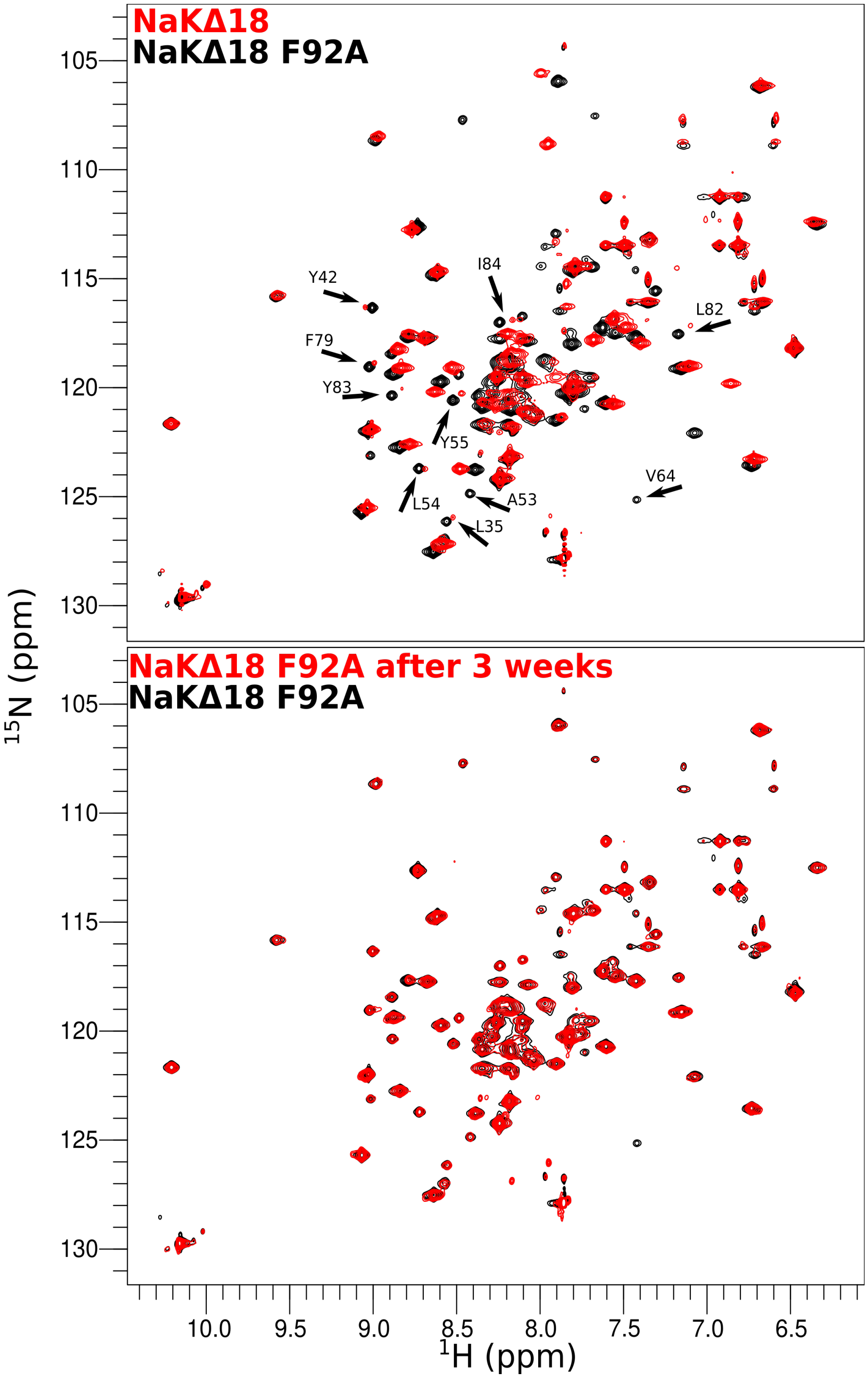


Figure SI11. Deuterium-hydrogen exchange of NaKΔ18 F92A as observed with $^{15}$N HSQC spectra. Overlay of wild-type NaKΔ18 (red) and NaKΔ18 F92A (black) in q=0.33 DMPC/DHPC bicelles upon initial purification (top). Residues which are usually poorly back-exchanged (Lewis2021) are indicated by arrows. Minimal change is observed over three weeks at 40 °C for NaKΔ18 F92A (bottom).

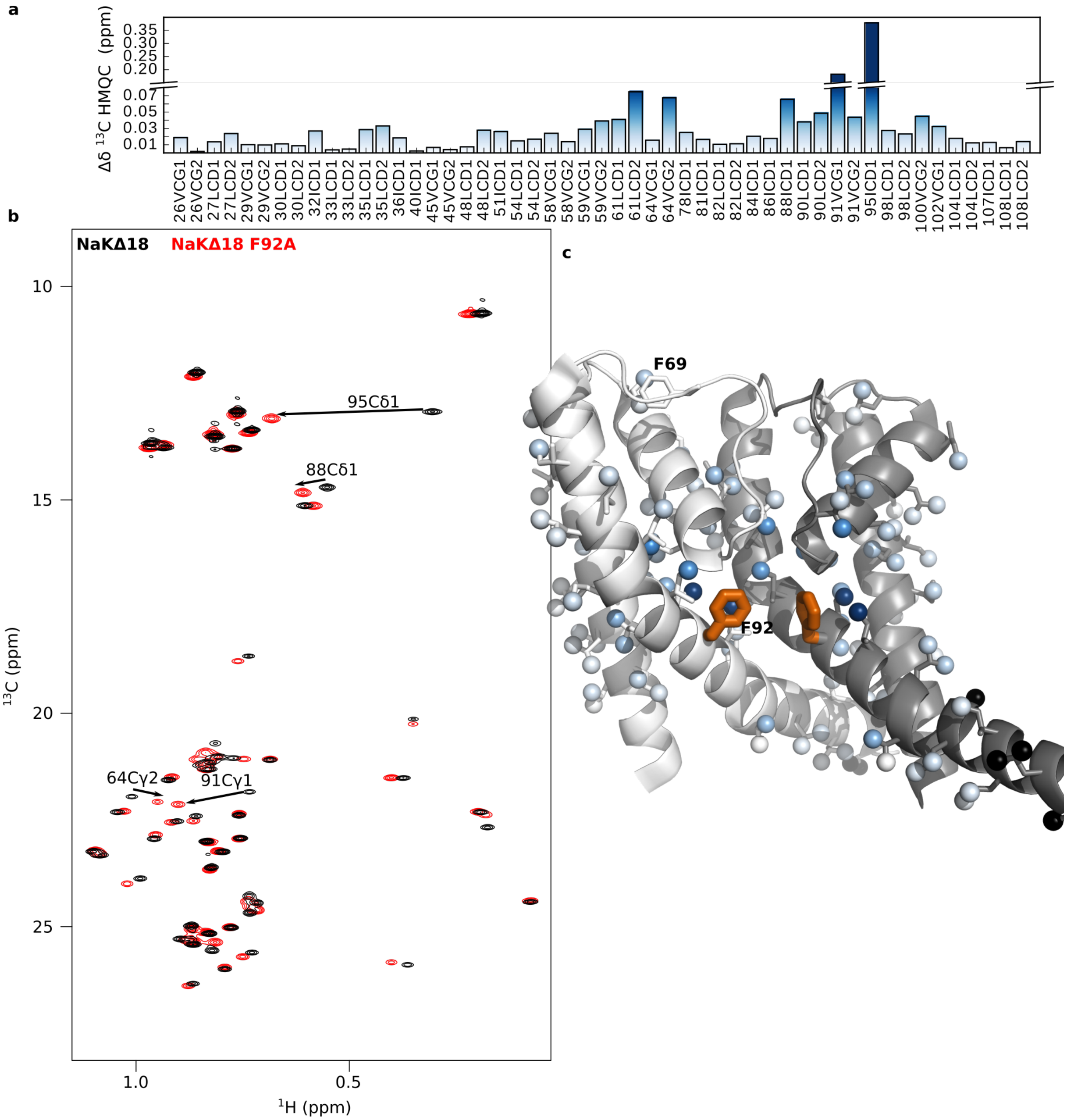


Figure SI12. Chemical shift difference between WT NaKΔ18 and NaKΔ18 F92A. $^{1}H$,$^{13}C$-HMQC spectra for NaK Δ18 (black) and F92A (red) reveal some significant chemical shift perturbation that are plotted as a bar graph (top) and on the methyl groups of the NaKΔ19 crystal structure (PDB 3E8H) using the same blue to white color scale. F92 is shown in thick orange sticks and F69 in white sticks. Black spheres indicate unassigned residues.

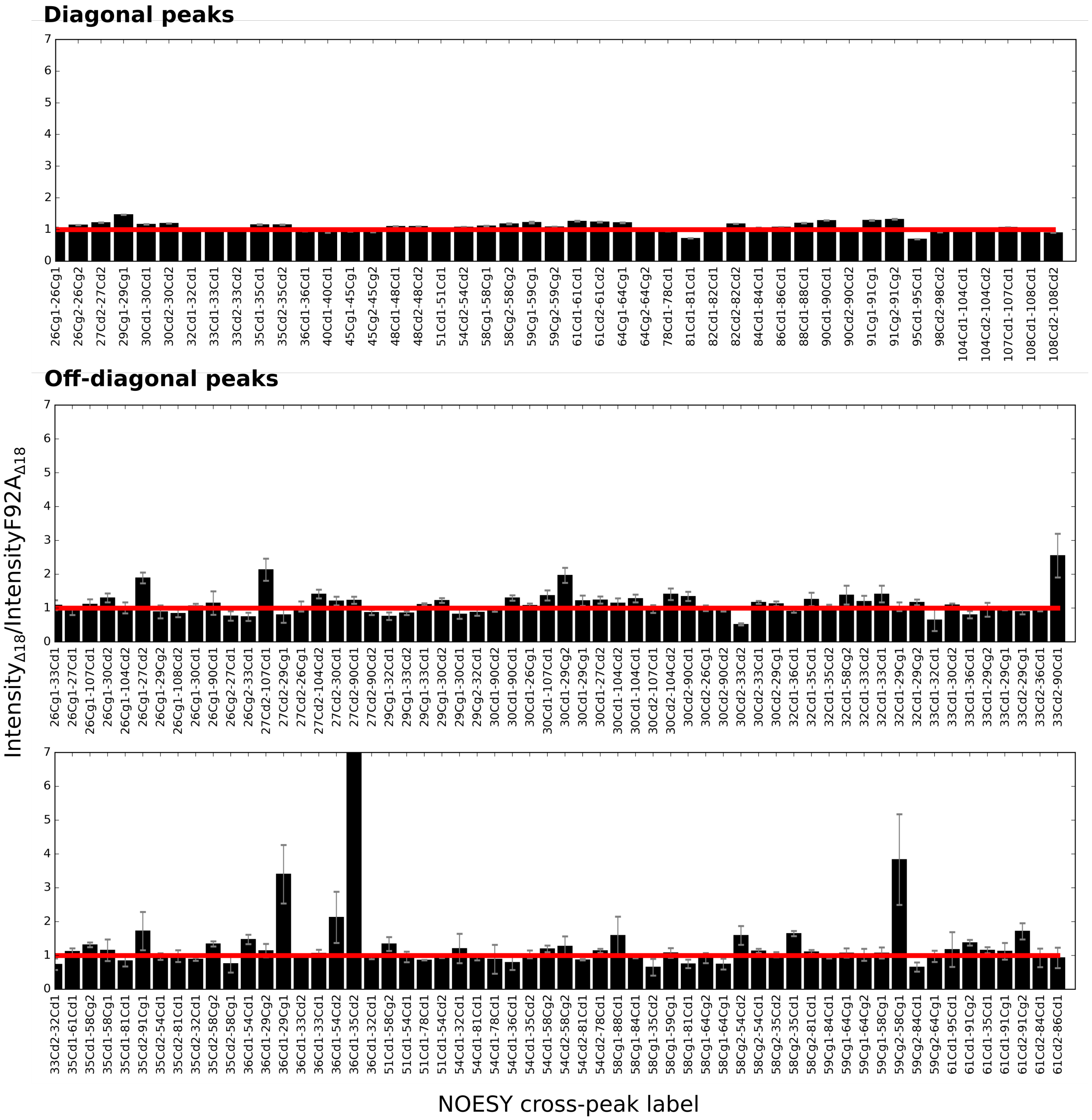


Figure SI13. Left: NaKΔ18 WT/NaKΔ18 F92A intensity ratios for all unambiguous off-diagonal and diagonal cross-peaks. Error bars represent propagated standard deviation of peak intensity. Cross-peak identity is displayed below the bars (name of resonance coming from diagonal peak first). Continued on the next page.

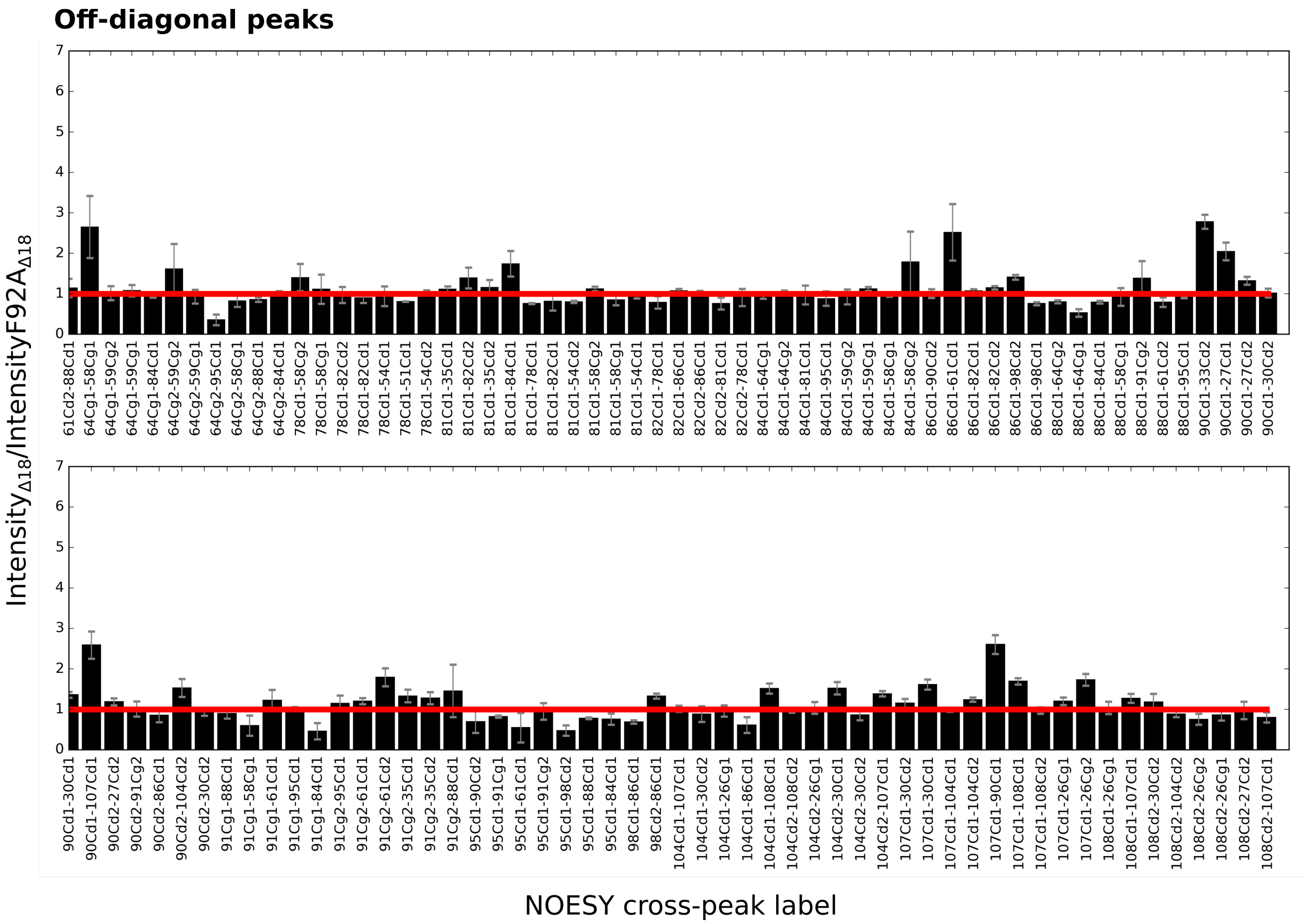


Figure SI13. Continued.

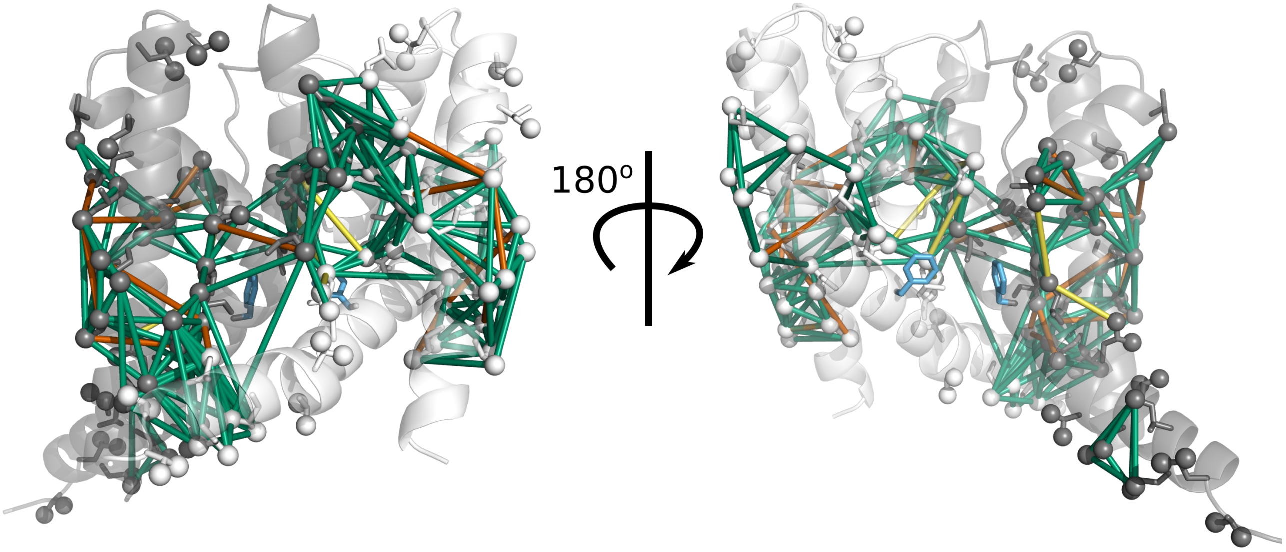


Figure SI14. The same data as in Figure SI12 displayed on the NaKΔ19 “open” structure (PDB 3E8H). Green lines represent ratios between 0.5 and 2. Orange lines indicate when the peak is at least 2x more intense in NaKΔ18, and yellow lines where the peaks is at least 2x less intense in NaKΔ18 as compared to NaK FL. Substituted F92 residue is blue.

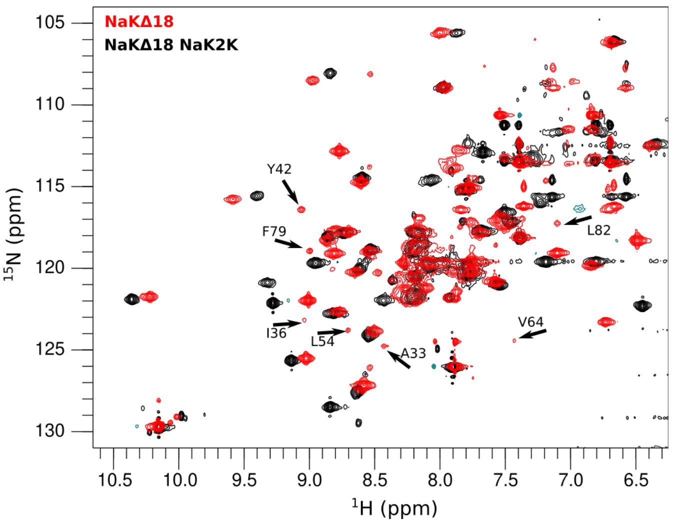


Figure SI15. Deuterium-hydrogen exchange of NaK2KΔ18. Overlay of WT NaKΔ18 (red) and NaK2KΔ18 (black) 1H-15N TROSY-HSQC spectra. Residues which are usually poorly back-exchanged are indicated by arrows.

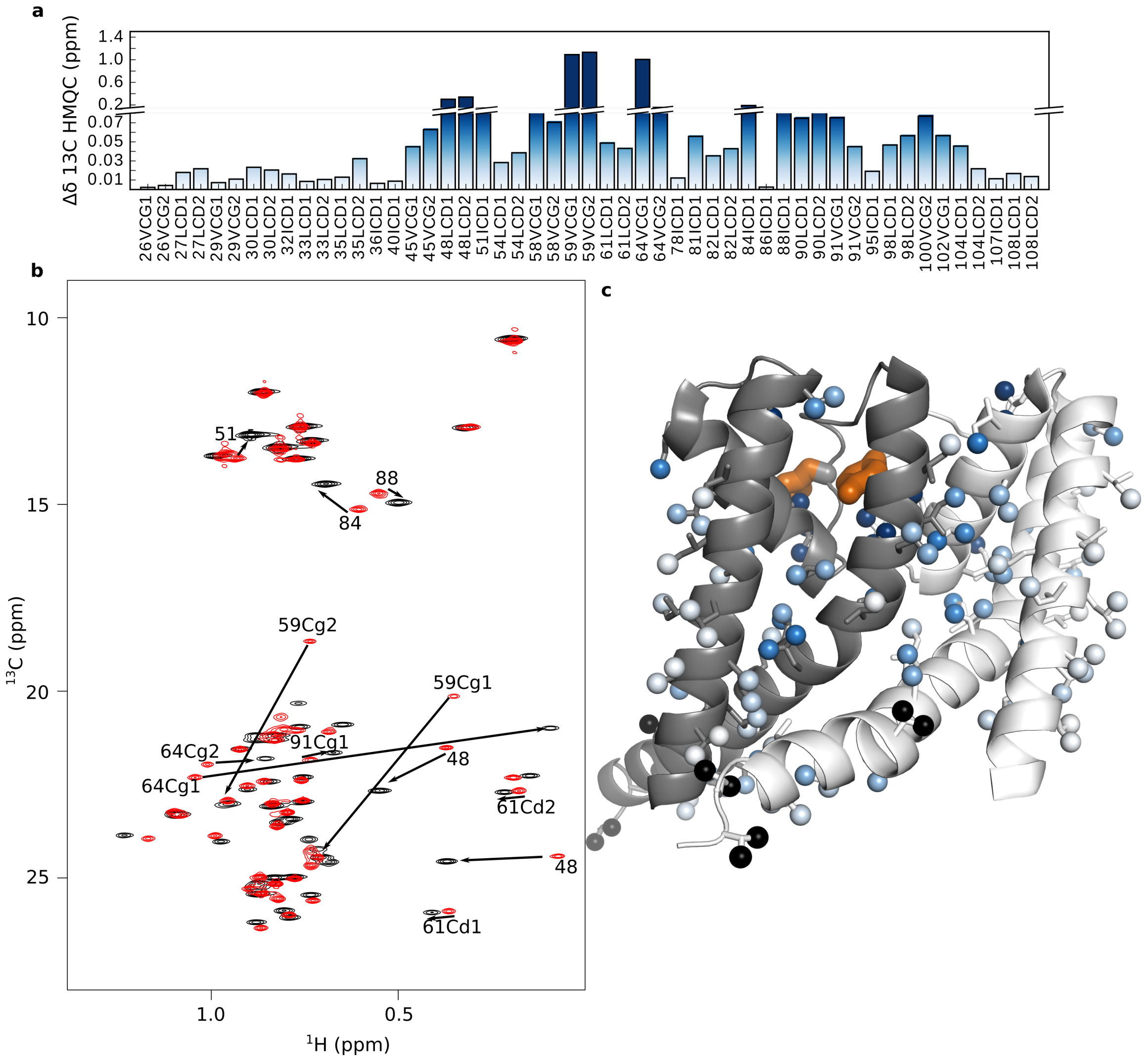


Figure SI16. Methyl chemical shift differences between WT NaKΔ18 and NaK2KΔ18. $^{1}$H,$^{13}$C-HMQC spectra of NaK Δ18 (black) and NaK2K (red) show some significant chemical shift perturbations (assignments as shown). The combined $^{1}$H,$^{13}$C chemical shift difference is shown as a bar graph (top) and plotted on the structure (PDB 3E8H, right) using the same white to blue color scale. Mutated residues (D66Y, N68D) are shown in thick orange sticks. Black spheres indicate unassigned residues.

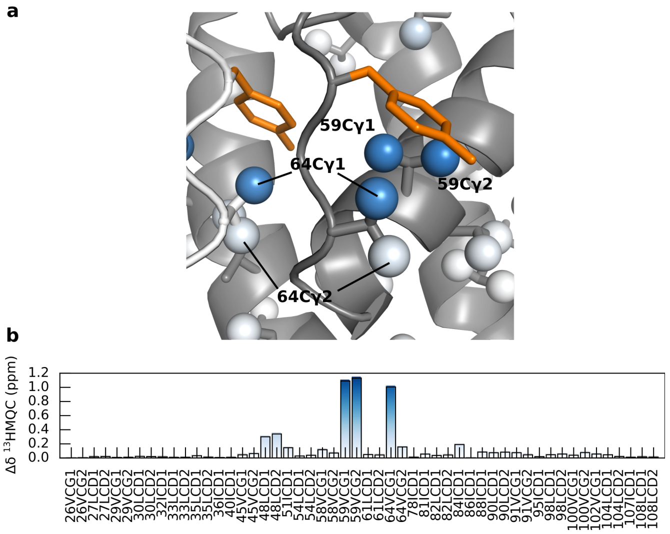


Figure SI17. a) structural representation of the aromatic ring of tyrosine introduced into NaK after D66Y mutation (orange) and b) methyl atoms undergoing very large CSPs relatively to the wild-type NaK spectra, as observed by NMR.

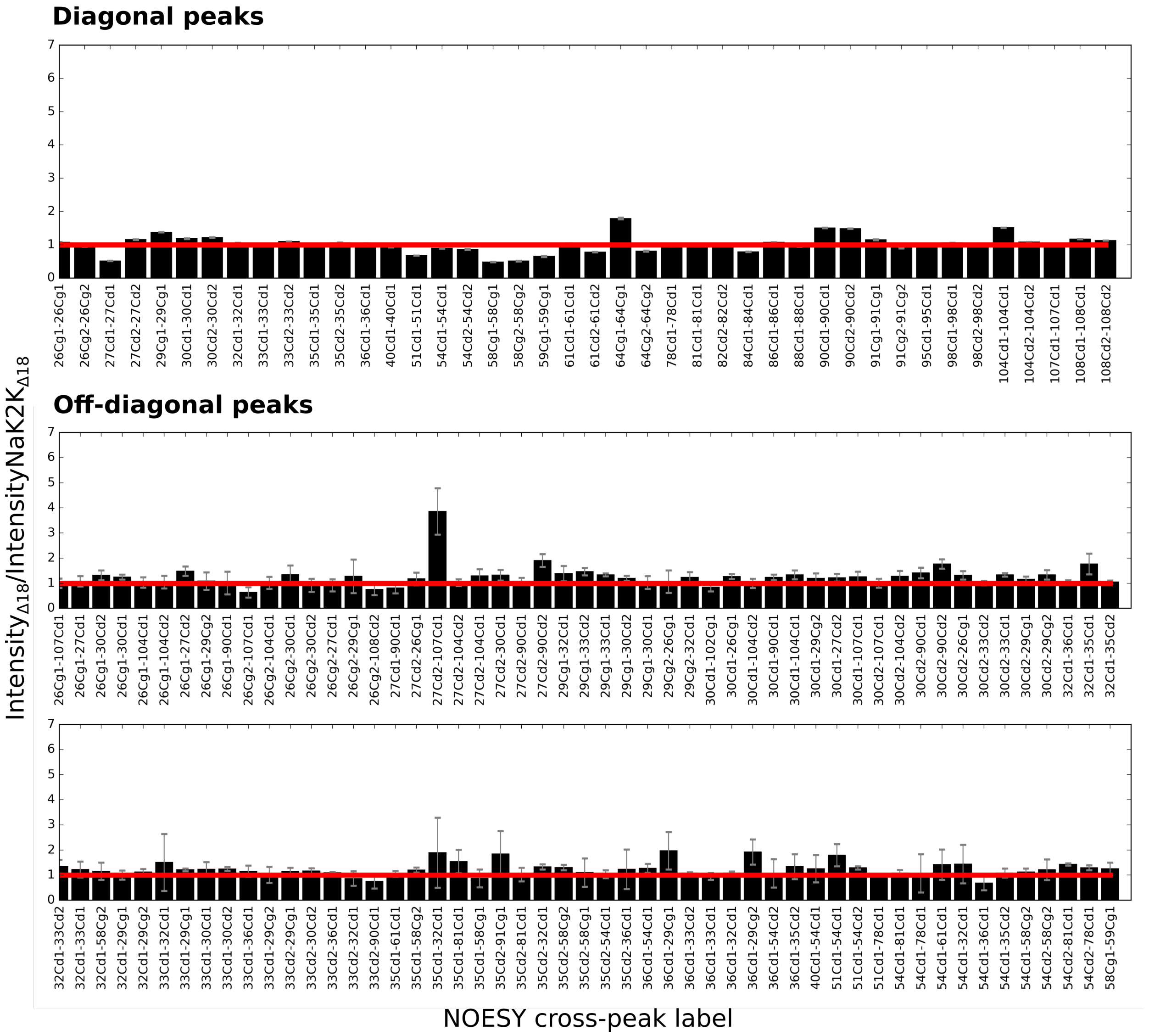


Figure SI18. NaKΔ18 WT/NaK2KΔ18 intensity ratios of all unambiguous off-diagonal and diagonal cross-peaks. Error bars represent propagated standard deviation of peak intensity. Cross-peak identity is displayed below the bars (name of resonance coming from diagonal peak first). Continued on the next page.

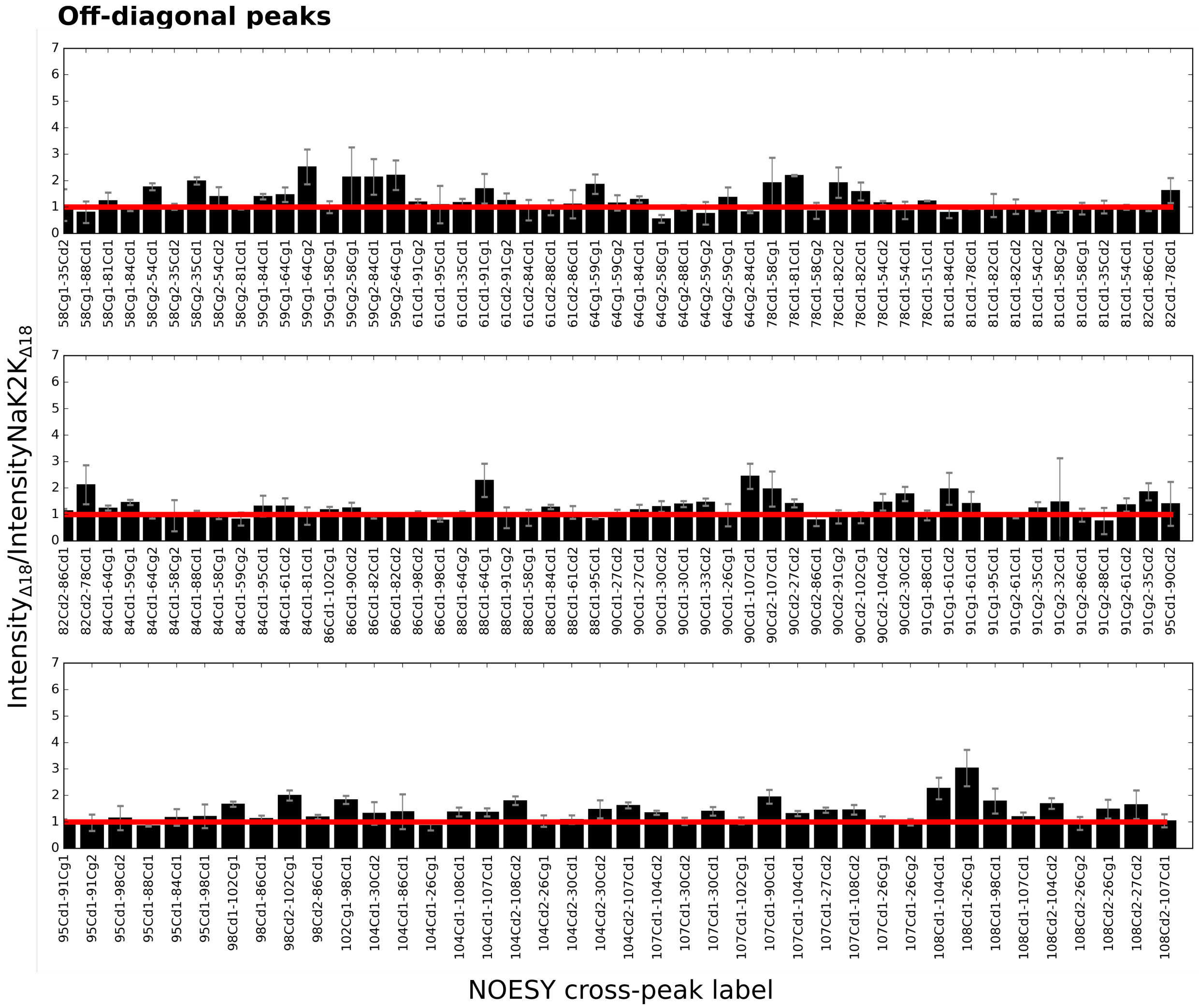


Figure SI18. Continued.

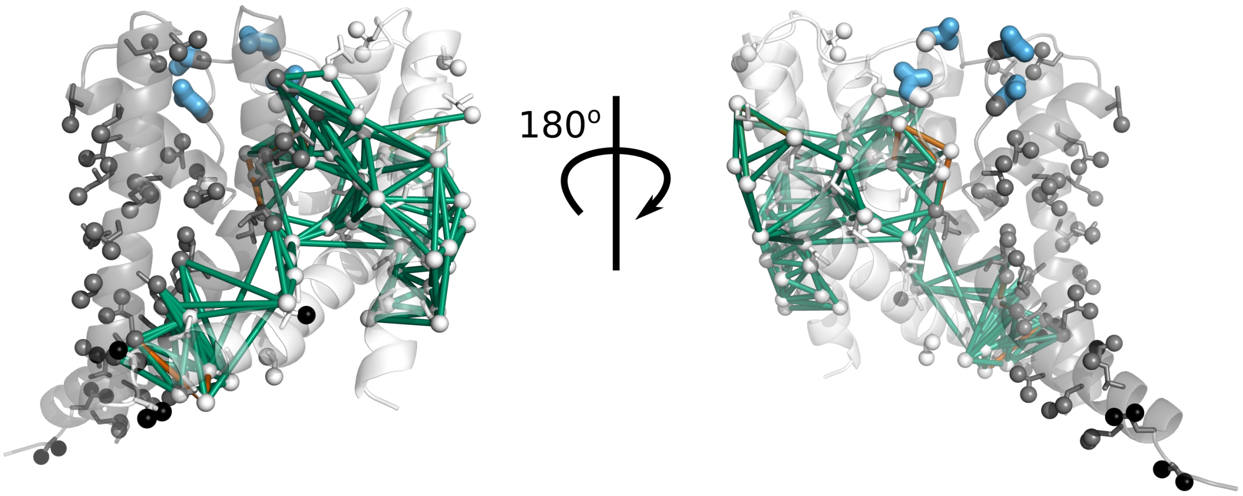


Figure SI19. The same data as in Figure SI18 is plotted on PDB 3E8H. Green lines represent ratios between 0.5 and 2. Orange lines indicate when the peak is at least 2x more intense in NaKΔ18, and orange lines where the peaks at least 2x less intense in NaKΔ18 as compared to NaK2K. Substituted residues are blue.
